# Unambiguous detection of SARS-CoV-2 subgenomic mRNAs with single cell RNA sequencing

**DOI:** 10.1101/2021.11.22.469642

**Authors:** Phillip Cohen, Emma J DeGrace, Oded Danziger, Roosheel S Patel, Erika A Barrall, Tesia Bobrowski, Thomas Kehrer, Anastasija Cupic, Lisa Miorin, Adolfo García-Sastre, Brad R Rosenberg

**Affiliations:** Department of Microbiology, Icahn School of Medicine at Mount Sinai, New York, NY 10035

**Author notes:** Address correspondence to Brad R Rosenberg.

## Abstract

Single cell RNA sequencing (scRNA-Seq) studies have provided critical insight into the pathogenesis of Severe Acute Respiratory Syndrome CoronaVirus 2 (SARS-CoV-2), the causative agent of COronaVIrus Disease 2019 (COVID-19). scRNA-Seq workflows are generally designed for the detection and quantification of eukaryotic host mRNAs and not viral RNAs. Here, we compare different scRNA-Seq methods for their ability to quantify and detect SARS-CoV-2 RNAs with a focus on subgenomic mRNAs (sgmRNAs). We present a data processing strategy, single cell CoronaVirus sequencing (scCoVseq), which quantifies reads unambiguously assigned to sgmRNAs or genomic RNA (gRNA). Compared to standard 10X Genomics Chromium Next GEM Single Cell 3′ (10X 3′) and Chromium Next GEM Single Cell V(D)J (10X 5′) sequencing, we find that 10X 5′ with an extended read 1 (R1) sequencing strategy maximizes the detection of sgmRNAs by increasing the number of unambiguous reads spanning leader-sgmRNA junction sites. Using this method, we show that viral gene expression is highly correlated across cells suggesting a relatively consistent proportion of viral sgmRNA production throughout infection. Our method allows for quantification of coronavirus sgmRNA expression at single-cell resolution, and thereby supports high resolution studies of the dynamics of coronavirus RNA synthesis.

## Introduction

Severe Acute Respiratory Syndrome CoronaVirus 2 (SARS-CoV-2) is the causative agent of COronaVIrus Disease-2019 (COVID-19), which as of January 2023 has caused over 663 million cases and over 6.7 million deaths globally(1, 2). Global efforts to understand the pathogenesis of SARS-CoV-2 infection have led to the development of vaccines and antiviral drugs, which have significantly reduced morbidity and mortality(3). “Omics” methods have been instrumental in studying SARS-CoV-2 in part because they have generated large amounts of data regarding host-viral interactions at unprecedented speed(4–15). Single-cell RNA sequencing (scRNA-Seq) studies in particular have been used to study multiple aspects of SARS-CoV-2 infection including but not limited to: viral tropism(16–22), peripheral immune changes(23–33), transcriptional changes induced by infection(34, 35), and to develop cell atlases of COVID-19 pathology(23, 24, 36, 37). Of note, most scRNA-Seq workflows have been developed and optimized for studies of eukaryotic transcription but not viral, specifically SARS-CoV-2, transcription. The performance of different scRNA-Seq methods to detect and quantify viral RNAs may impact the analysis and interpretation of such studies.

SARS-CoV-2 is a betacoronavirus with a 29 kB positive-sense, single stranded RNA genome(38, 39). SARS-CoV-2 generates genomic RNA (gRNA), subgenomic mRNAs (sgmRNAs), and negative-sense antigenomic RNA during active infection(40, 41). Both gRNA and sgmRNAs are polyadenylated, which enables detection by scRNA-Seq protocols that rely on poly-T primed reverse transcription(39–41). Translation of gRNA results in the production of one of two polyproteins, pp1a and pp1ab, which are subsequently cleaved into an array of non-structural proteins involved in pathogenesis and replication(39, 41). Translation of sgmRNAs generates structural and accessory viral proteins critical for virion production and pathogenesis(39, 41). Specific detection of sgmRNAs is therefore necessary for the analysis of viral gene expression dynamics within and across cells and viruses.

sgmRNAs are generated by discontinuous transcription events during negative strand synthesis(40). Transcription Regulatory Sequences (TRS), present in the 5′ leader sequence of the virus (TRS-L) and upstream of each ORF body (TRS-B), regulate this process(40). Template switching of the viral polymerase from a TRS-B to a TRS-L generates sgmRNAs with the 5′ leader sequence fused to the sgmRNA ORF body (**Figure 1A**)(40). These “nested” sgmRNAs share the viral ORF sequence downstream of the junction site in addition to a common leader sequence upstream of the junction site(40). This redundancy poses a challenge for standard scRNA-Seq data processing pipelines because reads mapping to redundant sgmRNA sequences are categorized as “ambiguous” and typically excluded from quantification. This problem has been addressed in bulk RNA-Seq by quantifying SARS-CoV-2 reads spanning leader-ORF junctions, which unambiguously identify sgmRNAs(12, 15). However, many scRNA-Seq methods do not sequence this region of sgmRNAs at significant coverage due to differences in library format and configuration of sequencing reads.

**Figure 1:**
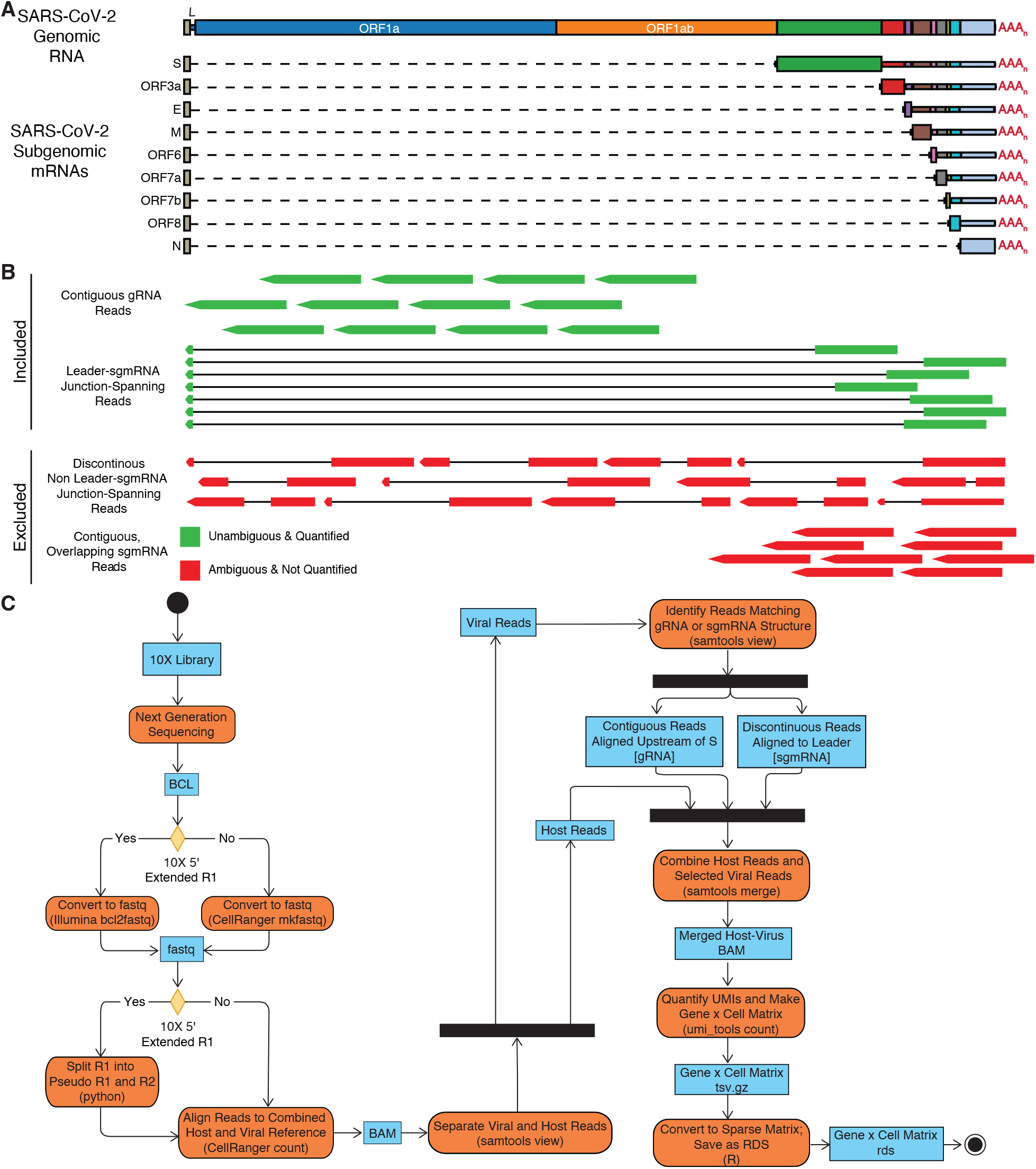
**A**. Illustration of SARS-CoV-2 genomic RNA, gRNA, and subgenomic RNAs, sgmRNAs. **B**. *Top:* Reads included for analysis by scCoVseq. Either: 1) contiguous reads mapping to ORF1a/b and therefore derived from gRNA or 2) discontinuous reads spanning the leader region and ORFS transcribed by sgmRNAs *Bottom*: Reads excluded from analysis by scCoVseq. Either: 1) discontinuous reads that do not include sequence mapping to the leader region or 2) contiguous reads that map to ORFs other than ORF1a/b, which are ambiguous. **C**. Activity diagram of scCoVseq pipeline. Blue rectangles indicate inputs/outputs for each stage. Orange rounded rectangles indicate a process in bold with software indicated.

We hypothesized that both experimental (i.e. scRNA-Seq library construction and sequencing strategies) and data processing decisions influence the ability to detect, resolve, and quantify SARS-CoV-2 RNA species with scRNA-Seq. We developed a data processing workflow, single cell coronavirus sequencing (scCoVseq), to quantify only unambiguous SARS-CoV-2 sgmRNA reads in scRNA-Seq data. We found that SARS-CoV-2 RNA detection differed by 10X Genomics Chromium scRNA-Seq library construction methods, due in part to ambiguity of the library fragments generated by each method. We show that 10X Chromium Next GEM Single Cell V(D)J (10X 5′) scRNA-Seq with an extended read 1 (R1) sequencing strategy maximized unambiguous SARS-CoV-2 reads and thereby increased detection of SARS-CoV-2 RNAs. Our method yielded similar sgmRNA quantities in aggregated scRNA-Seq data as detected with previously published bulk RNA-Seq methods(42). Using this method, we show that viral gene expression is highly correlated across infected cells, which may suggest that the relative proportion of viral sgmRNA expression is maintained throughout infection.

## Materials and Methods

### Cell lines and Viral Infection

Vero-E6 cells (ATCC, CRL-1586) were maintained in Dulbecco’s Modified Eagle Medium (DMEM, Corning) supplemented with 10% fetal-bovine serum (FBS) and 1% Penicillin Streptomycin (PSN, Fisher scientific), and routinely cultured at 37° C with 5% CO_2_. A549-ACE2 (previously described in (43, 44) were maintained in Dulbecco’s modified Eagle’s medium (Corning) supplemented with 10% fetal bovine serum (Peak Serum) and penicillin/streptomycin (Corning) at 37 °C and 5% CO2. All cell lines used in this study were regularly screened for Mycoplasma contamination with the MycoAlert Detection Kit (Lonza).

All SARS-CoV-2 propagations and experiments were performed in a Biosafety Level 3 facility in compliance with institutional protocols and federal guidelines. For experiments involving Vero E6 cells, SARS-CoV-2 (isolate USA-WA1/2020, BEI resource NR-52281) and control media (mock infected) stocks were grown by inoculating a confluent T175 flask of Vero-E6 cells (passage 2). Mock and SARS-CoV-2 infected cultures were maintained in reduced serum DMEM (2% FBS) for 72 hours, after which culture media was collected and filtered by centrifugation (8000 x g, 15 minutes) using an Amicon Ultra-15 filter unit with a 100KDa cutoff filter (Millipore # UFC910024). Concentrated stocks in reduced-serum media (2% FBS), supplemented with 50mM HEPES buffer (Gibco) were stored at -80°C. Viral titers were determined by plaque assay as previously described(43).

For experiments involving A549-ACE2 cells, a recombinant SARS-CoV-2 (rSARS-CoV-2) virus, rSARS-CoV-2 ORF6 M58R, was used as described elsewhere(45). All viral stocks were grown in Vero E6 cells as previously described and validated by genome sequencing (35). Sequencing was either performed using the MinION sequencer (Oxford Nanopore Technologies) or with the Nextera XT DNA Sample Preparation kit (Illumina) as described elsewhere (46, 47).

### scRNA-Seq

For scRNA-Seq experiments with Vero-E6 cells, cultures in 6 well plates were infected with SARS-CoV-2 at a MOI of 0.1, or with an equivalent volume of control media, in reduced-serum media (2% FBS) for 24 hours. To prepare cells for scRNA-Seq, mock and SARS-CoV-2 infected cultures were washed with calcium/magnesium-free PBS and disassociated using TrypLE (Gibco, 5 minutes at 37° C), after which samples were centrifuged (200 x g, 5 minutes), resuspended in calcium/magnesium-free PBS supplemented with 1% BSA, and counted. Mock and SARS-CoV-2 infected cell culture samples were filtered through a 40 μm FlowMi strainer (ScienceWare) and counted prior to loading on the 10X Genomics Chromium Controller according to manufacturer’s protocol. Mock and infected samples were loaded on separate lanes of a 10X Genomics Chromium Controller for either NextGEM Single Cell 3′ v3.1 (10X 3′), or NextGEM Single Cell V(D)J v1.1, (10X 5′).

scRNA-Seq gene expression libraries were prepared by 10X 3′ and 5′ library construction methods according to manufacturer’s guidelines. Final 10X 3′ mock and infected gene expression libraries and the 10X 5′ infected gene expression library were PCR amplified for 16 cycles while the 10X 5′ mock gene expression library was amplified for 14 cycles. 10X 3′ gene expression libraries were pooled and sequenced by short-read sequencing on an Illumina NextSeq 500 using a high output 150 cycle reaction kit according to manufacturers’ protocol with the following read lengths: read 1 28 nt; i7 index 8 nt; and read 2 130 nt. 10X 5′ gene expression libraries were also pooled and sequenced with 10X recommended read lengths (read 1 26 nt; i7 index 8 nt; and read 2 132 nt) or with extended R1 protocol (read 1 158 nt; i7 index 8 nt; no read 2).

For scRNA-Seq experiments with A549 cells, cultures were incubated with virus at an MOI of 2 in Dulbecco’s modified Eagle’s medium (Corning) supplemented with 2% fetal bovine serum (Peak Serum), 1% non-essential amino acids (Gibco), 1% HEPES (Gibco) and 1% penicillin/streptomycin (Corning) at 37 °C and 5% CO2. Cultures were incubated for 4 hours before supernatants were removed and replenished with fresh media. At 24 hours post-infection, cells were detached with TrypLE (Gibco), washed with PBS, and passed through a 70 μm filter. Finally, samples were processed using the Next GEM Single Cell Immune Profiling Assay (5’ RNA) single index kit (10X Genomics) according to the manufacturer’s instructions and as described previously, using 12 cycles of amplification during the cDNA amplification step (35). Sequencing was performed by the Icahn School of Medicine at Mount Sinai Genomics Core Facility using an Illumina NovaSeq with read lengths set to 200 bp for read 1, 8 bp for index 1, and 100 bp for read 2. Only read 1 was used for data denoted as 10X 5’ extended R1, and for analyses involving 10X 5’ data (i.e. not “extended”) R1 was truncated to 26 bases, the length required for cell barcode and UMI quantification, and read 2 was used without modification.

### scRNA-Seq Data Pre-Processing

#### Conversion of Illumina BCL files to fastq

Fastq files for standard sequencing of 3′ and 5′ gene expression libraries were generated using the mkfastq command in cellranger v.3.1.0 (10X Genomics). Fastq files for 5′ libraries sequenced with the extended R1 strategy were generated using bcl2fastq v2.20.0 (Illumina, Inc). Extended R1 fastqs were then separated into “pseudo R1” fastqs, containing the cell barcode and UMI, and “pseudo read 2 (R2)” fastqs, containing cDNA sequence, using a customized Python (v3.7.3) script (available at GitHub *link pending*) as follows.

The cell barcode and UMI are selected from the first 26 bp of R1. The subsequent 13 bp are derived from the template switch oligonucleotide and are ignored. The remaining nucleotides (and corresponding quality scores) are reverse complemented and stored as pseudo R2. The read header of the pseudo R2 fastqs are modified to reflect the format for standard R2 fastqs.

#### Downsampling fastqs to control for sequencing depth

To control for differences in sequencing depth for each Vero E6 library, read depth per library was downsampled to approximately 50,000 reads per cell. To generate a whitelist of cell barcodes for downsampling while accounting for transcriptional shutdown in SARS-CoV-2 infected cells (35), we generated preliminary gene x cell matrices for our dataset using cellranger/3.1.0 count (10X Genomics, Inc) to quantify and align reads to a host reference (African Green Monkey, ChlSab1.1) combined with SARS-CoV-2 transcripts as annotated by NCBI SARS-CoV-2 reference (NC_045512.2) with modifications for USA/WA01 strain for each dataset. The resulting output was analyzed in R (v4.0.4) with Seurat (v4.0.1)(48–50) to filter out putative doublets and empty droplets according to total UMIs/cell, number of genes/cell, and percent of mitochondrial gene expression. After filtering, putative cell-containing cell barcodes were output to a whitelist per library. Based on the these whitelists, the initial fastq files were downsampled using seqtk (v1.2)(51) to a total read depth of 50,000 multiplied by the number of cells in the library.

#### Preparation of empirically derived SARS-CoV-2 genome reference

Downsampled (for Vero E6 samples) and non-downsampled (for A549-ACE2 samples) fastq files were then mapped using cellranger count/3.1.0 (10X Genomics, Inc) to an empirically defined reference of SARS-CoV-2 sgmRNAs derived from previously reported SARS-CoV-2 (BetaCoV/Korea/KCDC03/2020) RNAs sequenced with long-read direct RNA Nanopore sequencing(12). These were downloaded from the UCSC Genome Browser Table Browser(52) after filtering for TRS-dependent transcripts and score > 900 and exporting to GTF format. Transcripts for previously unknown ORFs were excluded from the annotation. An additional annotation for “genomic RNA” was included which covered the entire length of the SARS-CoV-2 genome. Aligning the BetaCoV/Korea/KCDC03/2020 genome with USA/WA-CDC-WA1/2020 genome showed that the USA/WA-CDC-WA1/2020 reference had an additional 21 3′ adenosine nucleotides annotated. To account for this in our reference, we extended any annotations from BetaCoV/Korea/KCDC03/2020 that ended at the 3′ end of the genome by an additional 21 bases. This SARS-CoV-2 reference was appended to the host ChlSab1.1 Ensembl reference for Vero E6 analyses or GRCh38 version 100 human reference for ACE2-A549 analyses.

### scCoVseq

To unambiguously assign and quantify scRNA-Seq reads to SARS-CoV-2 RNAs, the cellranger output BAM was filtered for reads mapping to SARS-CoV-2 or ChlSab1.1 references using samtools (version 1.11)(53). SARS-CoV-2 aligned reads were then subset to likely genomic RNA reads or sgmRNA reads as follows. Genomic reads were defined as those containing no gaps in their alignment and mapping upstream of the start of the most 5′ sgmRNA, S. sgmRNA reads were defined as SARS-CoV-2 reads containing a gap and mapping in part to the 5′ leader sequence, defined as the 5′ proximal 80 nucleotides of the SARS-CoV-2 genome, and in part 3′ to the start of S. All other reads mapping to the SARS-CoV-2 genome were discarded. Reads passing these filtering steps were quantified with umi_tools (v1.0.0)(54). An R (v3.5.3) script using the Matrix (v1.2-18)(55) and readr (v1.3.1)(56) packages was used to convert this to a sparse matrix and save as an rds file. UMIs assigned to multiple genes were removed from the resulting matrix during downstream analysis.

### scRNA-Seq Data Analysis

#### Sashimi Plots

Reads from Vero E6 samples for 10X 3′, 10X 5′, and 10X 5′ extended R1 data that aligned to the SARS-CoV-2 reference by cellranger were subset from the cellranger output BAM file. Each BAM file was downsampled to approximately 1 × 10^6^ reads to control for differences in sequencing depth across libraries. Sashimi plots were generated with ggsashimi (v1.0.0)(57).

#### Classification of SARS-CoV-2 Infected Cells

scCoVseq-derived gene by cell matrices were loaded into R (v4.0.4) and analyzed with the Seurat (v4.0.1)(48–50) package. For each 10X method, mock and infected gene x cell matrices were merged with the Seurat merge command. To identify infected and bystander cells within SARS-CoV-2 treated cultures, euclidean distance between the z-scaled expression of SARS-CoV-2 sgmRNA UMIs per cell was clustered using k-medoids by the pam algorithm (k = 2) implemented in the cluster (v2.1.2) package(58). Output clusters were then compared for viral UMI expression per cell, and the cluster with more viral UMIs was classified as infected and the other as uninfected.

#### Comparison of SARS-CoV-2 RNA UMIs per scRNA-Seq Method

To examine the distribution of SARS-CoV-2 UMIs per cell by scRNA-Seq method, the 25^th^ percentile of total UMIs was quantified for all infected cells from each 10X method. Any cells with fewer UMIs than the minimum 25^th^ percentile of all samples were discarded, and all cells were subsequently downsampled to this same number of total UMIs/cell using the Seurat SampleUMI command. Each dataset was randomly downsampled to the same number of infected cells to equalize for differences in cell numbers, and viral sgmRNA UMIs/cell were plotted by scRNA-Seq method.

#### SARS-CoV-2 Read Distribution by scRNA-Seq Method

SARS-CoV-2 reads were defined as genomic or subgenomic using scCoVseq. Reads aligning to the SARS-CoV-2 reference that were excluded from scCoVseq were classified as ambiguous. The number of genomic, subgenomic, or ambiguous reads per million SARS-CoV-2 reads was calculated and plotted for each scRNA-Seq method.

#### Differential Expression Analysis

To explore expression differences between infected, bystander, and mock cells, differential expression testing with edgeR (v3.32.1) was performed with modifications for scRNA-Seq as previously described(59, 60). Viral genes were excluded from analysis, and only host genes expressed in at least 10% of cells were tested. To account for differences in RNA content of infected cells due to virus-induced transcriptional shutdown, all cells were downsampled to the 25^th^ percentile of total UMIs of infected cells. Cells with fewer UMIs than the threshold were excluded from analysis. Differential gene expression was performed with edgeR using a generalized linear model quasi-likelihood F test adapted with a term for gene detection rate(59, 60). Genes with an absolute log_2_ fold change greater than or equal to 1 and false discovery rate less than 0.05 were considered significant. For KEGG enrichment analysis, pairwise tests between mock, bystander, and infected cells were performed. Differentially expressed genes with an absolute log_2_ fold change greater than or equal to 1 and false discovery rate less than 0.05 were considered significant and subject to KEGG enrichment analysis using the KEGG annotations for African Green Monkey as implemented in the edgeR function kegga. *Quantification of SARS-CoV-2 sgmRNA Junction Sites*. We explored the ability of our extended R1 sequencing to detect SARS-CoV-2 sgmRNA junctions using STARsolo (version 2.7.8a)(61). Aligned reads were re-mapped to the empirical SARS-CoV-2 annotation described above and junction sites per cell were quantified. The resulting junction per cell matrix was plotted in R (v4.0.4).

### Bulk RNA-Seq

#### Bulk RNA-Seq library preparation and sequencing

A549-ACE2 samples for bulk RNA sequencing were lysed in Trizol Reagent (Invitrogen) and total RNA was extracted using the miRNeasy mini kit (Qiagen) per the manufacturer’s instructions. DNAse treatment was performed on isolated RNA using the RNA Clean and Concentrator Kit (Zymo). Total RNA was examined for quantity and quality using the TapeStation (Agilent) and Quant-It RNA (ThermoFisher) systems. RNA samples with sufficient material (10 pg–10 ng) were passed to whole-transcriptome library preparation using the SMART-Seq v4 PLUS Kit (Takara Bio) following the manufacturer’s instructions. This library construction method was selected due to molecular similarities (e.g. 5′ template switch adaptor addition, full length cDNA amplification, etc.) to 10X Genomics Chromium scRNA-Seq library preparation. Briefly, total RNA inputs were normalized to 10ng in 10.5 µl going into preparation. 3′ ends of cDNA were then adenylated prior to ligation with adapters utilizing unique dual indices (96 UDIs) to barcode samples to allow for efficient pooling and high throughput sequencing. Libraries were enriched with PCR, with all samples undergoing 14 cycles of amplification prior to purification and pooling for sequencing. Bulk RNA sequencing was conducted on dual index libraries using a 300 cycle Mid Output kit on an Illumina NextSeq 500 with standard read configurations for R1, i7 index, i5 index, and R2:150, 8,8,150.

#### Bulk RNA-Seq analysis

Raw BCL files were converted to fastq files using bcl2fastq/2.20.0 (Illumina, Inc). For quantification of SARS-CoV-2 sgRNA and gRNA expression, the periscope/0.1.2 package was used with the technology argument set to “illumina” (42). Finally, sgRNA reads per total mapped reads were calculated.

### Flow Cytometry

Vero E6 cells were fixed with 4% paraformaldehyde at room temperature for a minimum of 24 hours, washed once with PBS and permeabilized with 1X perm-wash buffer (BD Biosciences) for 5 minutes. SARS-CoV nucleocapsid (N) antibody (clone 1C7C7) (kindly provided by Thomas Moran, Icahn School of Medicine at Mount Sinai, New York, NY), conjugated to AlexaFluor 647 was diluted 1:400 in perm-wash buffer, and added directly to samples. Samples were then incubated at room temperature for 40 minutes in the dark. After staining, samples were washed once with 1X perm-wash buffer, once with PBS, resuspended in FACS buffer (PBS supplemented with 1% FBS), and acquired on a Gallios flow cytometer (Beckman-Coulter). For all viral infections, analysis was performed with FlowJo software (v10.7.1, Becton Dickinson), excluding cell doublets and debris and gating according to mock infected populations.

### Immunofluorescence microscopy

Vero E6 were seeded in 6-well plates (Falcon) with one glass coverslip (Fisher Scientific) per well. After 24 hours post infection, cells were washed with PBS and fixed with 4% paraformaldehyde (Fisher Scientific) overnight. Fixed cells were permeabilized using 0.1% Triton-X (Fisher Scientific) in PBS and blocked with 4% bovine serum albumin (BSA, Fisher Scientific) in PBS. Blocked coverslips were incubated with SARS-N-1C7 (1:500 in 4% BSA PBS) overnight at 4C, washed three times with PBS, and incubated for 45 minutes with 1:500 AlexaFluor 488-conjugated anti-mouse (Invitrogen, 1:500 in 4% BSA PBS) plus DAPI (Thermo Fisher Scientific, 1:1000 in 4% BSA PBS) at room temperature. Coverslips were then stained with phalloidin (1:400 in PBS) for 1 hour at room temperature and washed again three times with PBS. Coverslips were mounted using Prolong Diamond (Life Technologies P36970). Confocal laser scanning microscopy was performed using a Leica SP5 DMI (ISMMS Microscopy CoRE and Advanced Bioimaging Center) with a 40X/1.25 oil objective. Images were collected at a resolution of 512 × 512 pixels in triplicate per slide. Images were processed and analyzed using LAS X and CellProfiler (v4) (62).

### Data Availability

Raw and processed scRNA-Seq data are available at NCBI GEO (accession GSE189900) and analysis code is available at GitHub (*link pending*).

## Results

SARS-CoV-2 generates gRNA and sgmRNAs during infection, which are highly redundant in their sequences (**Figure 1A**). In scRNA-Seq processing pipelines, reads mapping to redundant sequences are “assigned” to all genes containing that sequence and are typically excluded from quantification steps. We therefore identified read structures that could unambiguously identify gRNA or different species of sgmRNAs to allow for their specific quantification (**Figure 1B**). Reads derived from gRNA should be contiguous and could map anywhere on the SARS-CoV-2 genome. Reads derived from sgmRNA could be either gapped or contiguous and could map to the 5′ leader and/or downstream of the start site of S, the most 5′ sgmRNA. Because contiguous reads mapping downstream of S could derive either from gRNA or sgmRNAs, they were excluded from quantification. We therefore defined gRNA reads as contiguous reads aligning upstream of regions contained in sgmRNAs. sgmRNA reads were defined as discontinuous reads spanning the leader region and regions used by sgmRNAs. Reads that did not match either of these formats could not be unambiguously assigned to gRNA or a sgmRNA and were therefore excluded from quantification (**Figure 1B**). With this framework, we developed scCoVseq to quantify unambiguous genomic and subgenomic viral reads (**Figure 1C**). Using scCoVseq, we compared the abilities of different scRNA-Seq methods to quantify SARS-CoV-2 RNAs.

In the widely available Chromium scRNA-Seq method developed by 10X Genomics, Inc, there are two formats for droplet-based scRNA-Seq: 10X 3′ and 10X 5′. 10X 3′ generates library fragments derived from the 3′ regions of polyadenylated RNAs within a cell (**Figure 2A**). Because sgmRNAs share all viral sequence 3′ of the leader-body junction site, 10X 3′ library fragments derived from SARS-CoV-2 heavily cover the 3′ end of the viral genome and do not contain leader-ORF junctions (**Figure 2D**). These reads are limited in their ability to differentiate gRNA from sgmRNA or distinguish different sgmRNA species. 10X 5′ generates library fragments from the 5′ termini of polyadenylated RNAs (**Figure 2B**). These fragments are on average approximately 500 bp long (according to the manufacturer’s documentation) and would be expected to contain leader-ORF junctions of SARS-CoV-2 sgmRNAs. The transcript read (R2), however, derives from the 3′ end of these fragments and at the recommended read length of 91 bases is not long enough to consistently sequence into the leader-sgmRNA junction site (**Figure 2B**). Reads from 10X 5′ can therefore contain some but not all junctions (**Figure 2E**). We reasoned that we could use 10X 5′ library fragments to detect junction-spanning reads by sequencing from the 5′ end of the fragment. To do this we extended R1, which is normally used to sequence the cell barcode and UMI, to sequence into the leader-body junction site (**Figure 2C**). Using 10X 5′ with extended R1, we were able to sequence more leader-sgmRNA junction sites and increase our ability to unambiguously quantify sgmRNAs (**Figure 2F**). Indeed 10X 5′ extended R1 increased the number of leader-sgmRNA spanning reads over 10X 5′ and 10X 3′ (**Figure 2G**).

**Figure 2:**
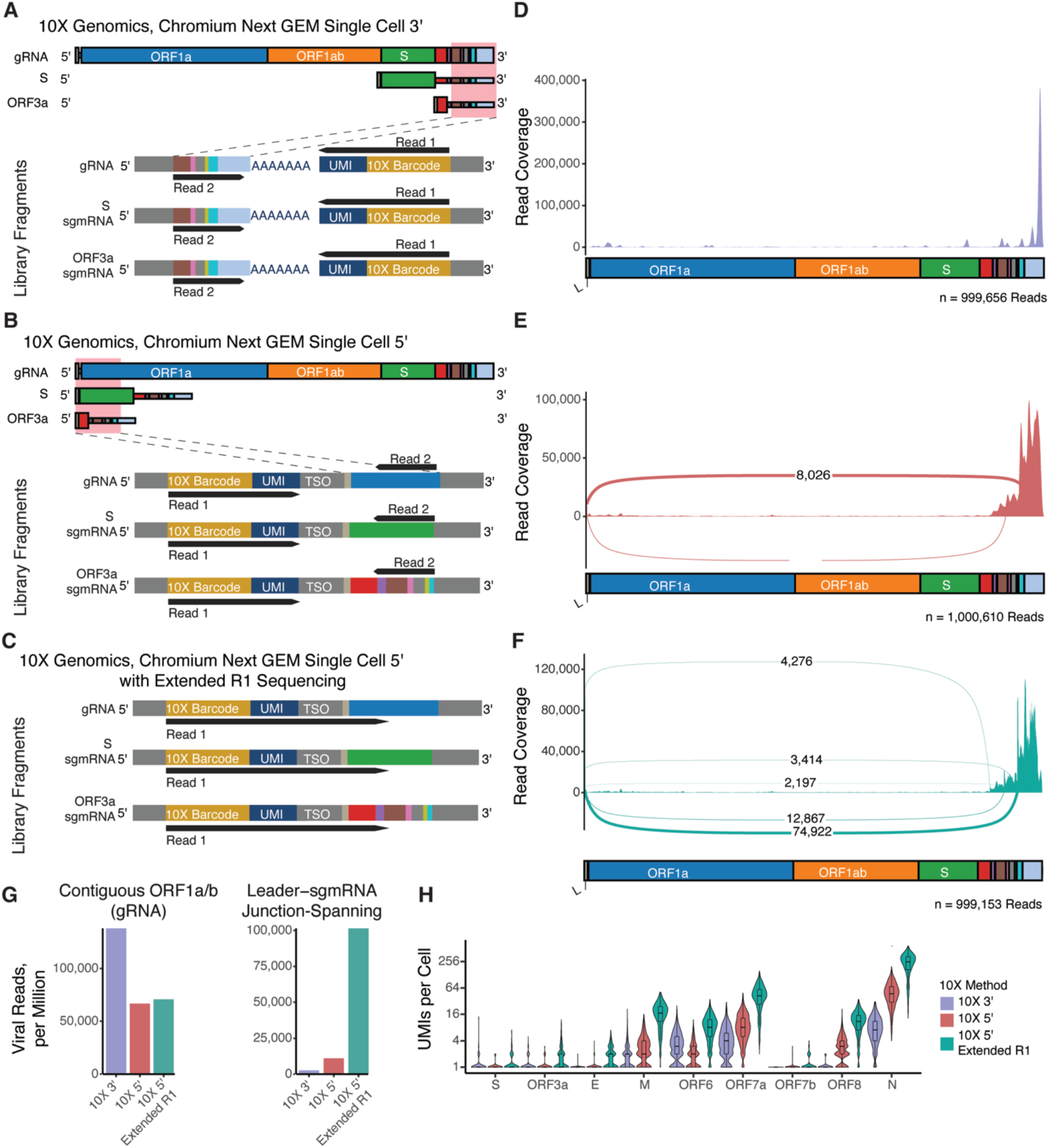
**A-C**. Illustration of gRNA and S and ORF3a sgmRNAs. Red box indicates regions contained in final 10X library. *Lower:* Example illustration of 10X library fragments derived from gRNA and S and ORF3a sgmRNAs with sequencing read 1 and read 2 indicated. 10X 3′ (**A**), 10X 5′ (**B**), and 10X 5′ extended R1 (**C**) libraries are illustrated. **D-F**. Sashimi plot of 10X 3′ (**D**), 10X 5′ (**E**), and 10X 5′ extended R1 (**F**) reads (Vero E6 cells) mapped to the SARS-CoV-2 genome filtered to show only junctions supported by at least 1,000 reads. Total number of reads visualized is indicated in the bottom right. **G**. Reads per million reads mapped to SARS-CoV-2 reads in 10X 3′, 10X 5′, or 10X 5′ with extended R1 data (Vero E6 cells). **H**. UMIs per cell for all sgmRNAs in infected cells in each dataset (Vero E6 cells). Each dataset was downsampled to an equal number of infected cells and each cells’ total UMIs were downsampled to the same value to control for differences in sequencing depth. The leader region is enlarged in illustrations of the genome for visibility. L = Leader.

Unexpectedly we observed a larger number of reads classified as gRNA in 10X 3′ compared to 10X 5′ or 10X 5′ with extended R1 (**Figure 2G**), and there appears to be increased read coverage in ORF1a and ORF1ab in the 10X 3′ library as well (**Figure 2D**). This phenomenon was also reported by Ravindra *et al* in their 10X 3′ libraries, who showed that these are non-canonical SARS-CoV-2 transcripts and not artifacts of the 10X 3′ library preparation (21). It is unclear at this time, however, why these transcripts may be better detected with 10X 3′ than 10X 5′.

When quantified with scCoVseq, we found that 10X 5′ extended R1 yields more UMIs for each sgmRNA per cell compared to 10X 5′ or 10X 3′ (**Figure 2H**). Importantly, the average host gene expression per sample was significantly correlated across 5′ methods, suggesting that host gene measurements were minimally affected by 10X 5′ extended R1 (**Supplemental Figure 1A-C**). For select host genes, we observed higher detection in 10X 5′ than 10X 5′ extended R1 in Vero cell data. Interestingly, however, when analyzing data from human A549 cells, we did not see any notable differences in quantification of host genes between 10X 5′ and 10X 5′ extended R1 (**Supplemental Figure 1D**). These differences, which are apparent for a small subset of total genes, may be a consequence of incomplete annotation of the African Green Monkey transcriptome reference, as they are not apparent with human data mapped to the more thoroughly curated human reference. Furthermore, in all of these analyses, viral genes were shown to be detected at higher levels (in aggregate) with 10X 5′ extended R1 without impacting host gene quantification. Taken together, 10X 5′ libraries sequenced with extended R1 sequencing results in a greater number of unambiguous reads derived from sgmRNAs over 10X 3′ or 10X 5′, and consequently recovers more sgmRNA UMIs/cell without affecting host gene quantification.

We next explored the ability of scCoVseq to quantify sgmRNAs as compared to previously published methods. To do this, we aggregated (to “pseudobulk” profiles) scRNA-Seq data of ACE2-expressing A549 cells infected with recombinant SARS-CoV-2 (rSARS-CoV-2) ORF6 M58R, an attenuated mutant that allows for high MOI infection without excessive cell death (45). In parallel, triplicate bulk RNA-Seq libraries were prepared from these cultures and quantified with periscope, a previously published method to quantify SARS-CoV-2 sgmRNAs in bulk RNAseq data using partial alignments of viral reads to TRS and ORF sequence(42). We compared the detection of sgmRNAs in aggregated “pseudobulk” scRNA-Seq data to the mean expression values in the bulk RNA-Seq data. We further compared 10X 5′ and 10X 5′ extended R1 data as well as data from each quantified with either CellRanger or our scCoVseq method (**Figure 3**). We found that either using 10X 5′ extended R1 sequencing or quantifying 10X 5′ with scCoVseq improved the correlation between aggregated scRNA-Seq data and bulk RNA-Seq data. The combination of 10X 5′ extended R1 and scCoVseq did not appear to improve further the correlation between scRNA-Seq and bulk RNA-Seq data. This analysis also showed that scRNA-Seq tends to detect higher levels of N and ORF6 sgmRNA and lower levels of ORF3a and M sgmRNA than bulk RNA-Seq. This may be due to differences in the properties of single cell as compared to bulk RNA-Seq library preparation or differences in the quantification of reads by the analysis pipelines used for each. Overall, these data suggests that 10X 5′ with extended R1 or scCoVseq improve the correlation of viral sgmRNA with bulk RNA-Seq quantification, but as described above, only the combination of both approaches maximizes the number of UMIs detected per sgmRNA per cell.

**Figure 3:**
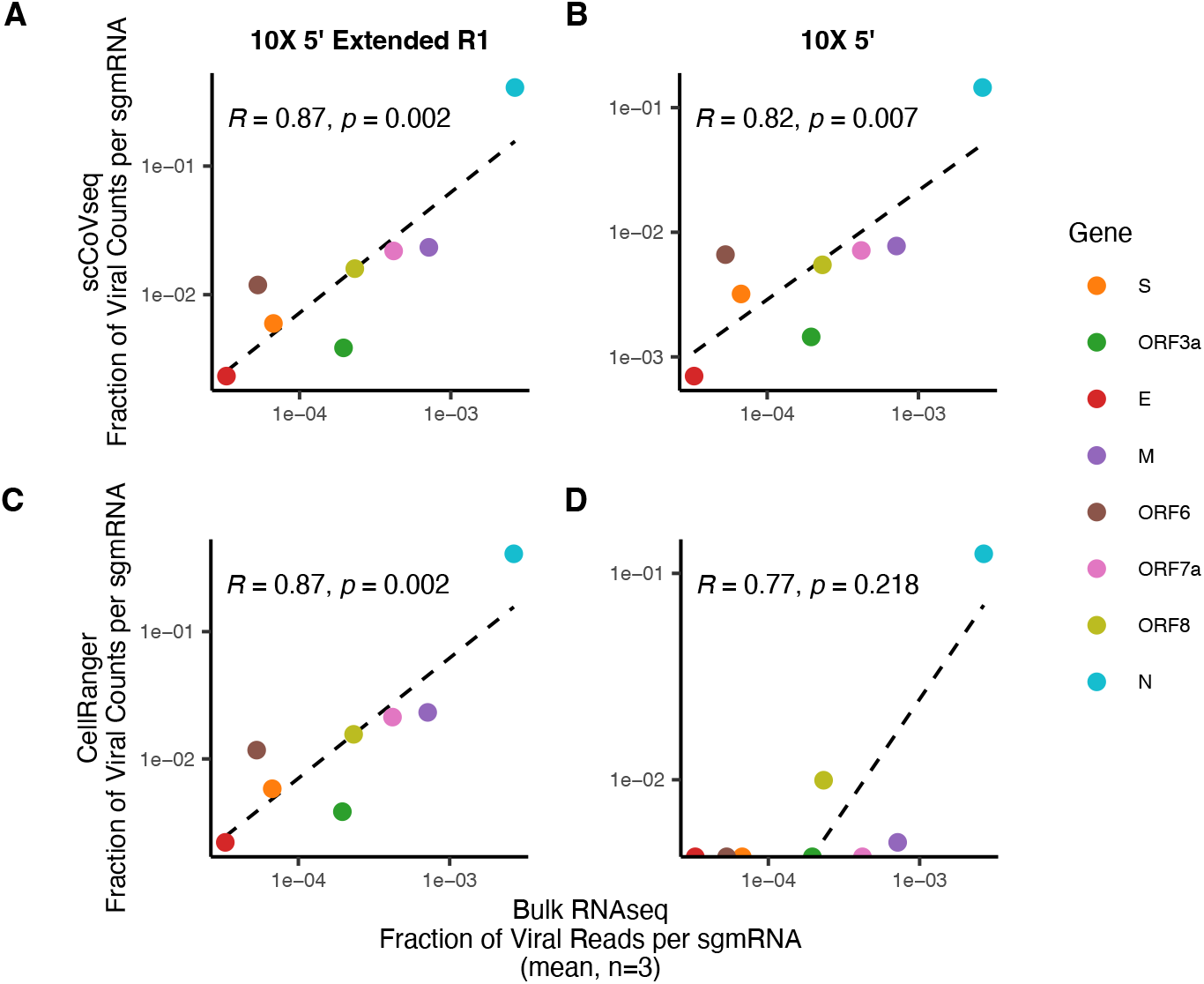
**A-C**. Correlation between viral sgmRNA quantification in bulk RNA-Seq by periscope and aggregated “pseudobulk” scRNA-Seq data of rSARS-CoV-2 ORF6 M58R infected ACE2-expressing A549 cells. The mean of three values for the fraction of total viral reads derived from each sgmRNA as quantified in bulk RNA-Seq are show on the x-axis. The fraction of total viral UMIs derived from each sgmRNA as quantified in scRNA-Seq are shown on the y-axis. Each point indicates a specific ORF. A regression line is fit to each graph, and Pearson’s correlation and resultant p values are shown for each analysis.

Next we sought to assess the relative expression of sgmRNAs during SARS-CoV-2 infection in single cells. We analyzed Vero E6 cells 24 hours post infection with SARS-CoV-2 at an MOI of 0.1 prepared with 10X 5′ extended R1 and analyzed with scCoVseq. We were able to quantify sgmRNAs and gRNA at single-cell resolution (**Figure 4A**). We classified infected cells using a k-medoid clustering approach based on sgmRNA expression (**Figure 4B**). We found that this classification method effectively separated cells with high levels of viral RNA from cells with low viral RNA (**Supplemental Figure 2B**) and detected a similar percentage of infected cells as detected using flow cytometry and immunofluorescence microscopy of the same cultures (**Supplemental Figure 3C**). We observed a progressive increase in all viral RNAs as total viral RNA UMI count increased in infected cells (**Figure 4C**). We further found that viral gene expression was highly correlated across cells, suggesting that the relative proportions of viral gene expression are tightly correlated throughout infection with SARS-CoV-2 (**Figure 4D**). These findings were also seen in rSARS-CoV-2 ORF6 M58R infected ACE2-expressing A549 cells (**Supplemental Figure 2A, C**). Importantly we showed that this trend was less clear in methods with suboptimal detection of SARS-CoV-2 RNAs, i.e., 10X 3′ and 10X 5′, which supports the utility of our method in studying viral gene expression at single-cell resolution (**Figure 4D**).

**Figure 4:**
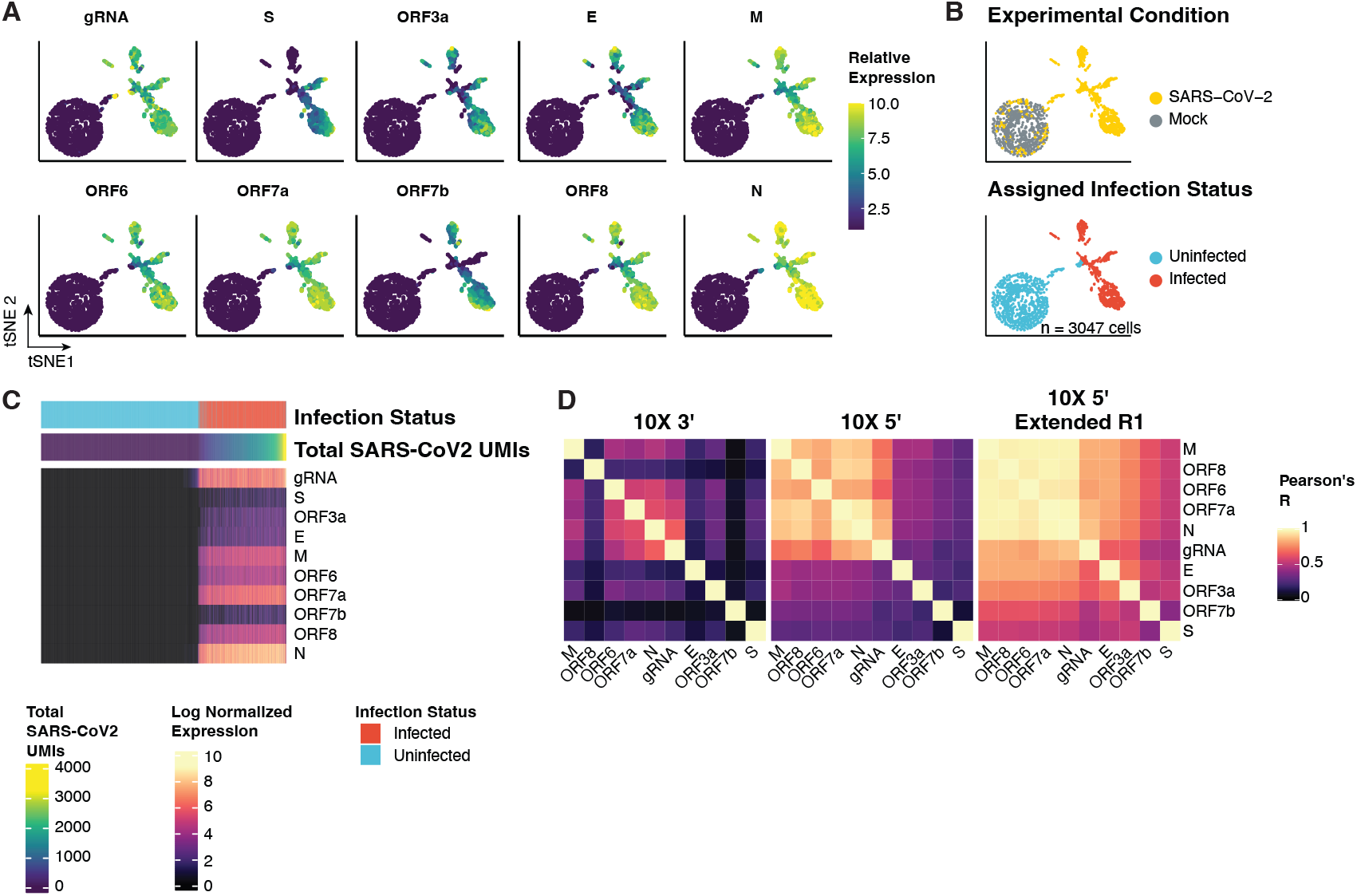
**A**. Viral gene expression in n = 3,047 mock and infected Vero E6 cells analyzed with 10X 5′ extended R1. Cells are embedded in tSNE space derived from Euclidean distance of scaled viral sgmRNA expression. **B**. Experimental condition and assigned infection status of cells, corresponding to tSNE in (A). **C**. Expression of viral genes in infected and uninfected Vero E6 cells. Cells are represented as columns with the expression of each viral gene indicated by color. Cells are ordered left to right by increasing total SARS-CoV-2 viral UMIs. Cells with total SARS-CoV-2 viral UMIs greater than the 99^th^ percentile are “clipped” to the 99^th^ percentile value for improved visibility. Cells are annotated above with assigned infection status. **D**. Correlation of viral gene expression across single cells as measured in 10X 3′, 10X 5′, and 10X 5′ extended R1.

Using our infection classification, we performed differential expression testing of infected Vero E6 cells compared to bystander cells within the same culture as well as to cells from a mock culture. As previously described (35), we observed suppression of many host genes in infected cells accompanied by an upregulation of cellular stress response genes (**Supplemental Figure 4A, B**). We further observed that, while bystander and mock cells had similar gene expression, a small number of genes were upregulated in bystander cells compared to mock cells. This is especially notable given the inability of Vero E6 cells to produce interferons in response to viral infection (63). KEGG enrichment analysis of differentially expressed genes in pairwise comparisons of infected, mock, and bystander cells showed that genes previously characterized as “related to COVID19” were enriched in our infected cells supporting our method for infection classification (**Supplemental Figure 4C**).

Finally, we explored the ability of our method to detect noncanonical leader-sgmRNA junctions in single cells. Multiple studies have reported on a variety of noncanonical sgmRNA structures detected in bulk RNAseq studies of SARS-CoV-2 infection including TRS-independent sgmRNAs (12, 13), but the physiological significance of these RNAs is currently incompletely understood. We hypothesized that 10X 5′ extended R1 might be able to detect junction sites corresponding to noncanonical sgmRNA structures because of its increased read coverage at expected junction sites. We observed a variety of noncanonical sgmRNA structures including TRS-independent sgmRNAs in our Vero E6 data sequenced with 10X 5′ extended R1 (**Supplemental Figure 5**). While our data were too sparse to study cell-type specific expression of sgmRNA junctions, future studies may be able to leverage 10X 5′ extended R1 to identify differential viral or host gene expression patterns associated with expression of certain sgmRNA junctions.

## Discussion

In this study, we examined the ability of two commonly used scRNA-Seq methods, 10X 3′ and 10X 5′, to detect and quantify SARS-CoV-2 derived RNAs with a focus on sgmRNAs. Because of the redundant nature of coronavirus sgmRNA sequences, we developed scCoVseq, which unambiguously quantifies both sgmRNAs and gRNAs in 10X data. We found that 10X methods detect unambiguous leader-sgmRNA junction-spanning reads to different degrees. We were able to increase the detection of leader-sgmRNA junction-spanning reads by extending the length of R1 during sequencing of 10X 5′ libraries, an approach we term 10X 5′ extended R1 sequencing. We further showed that analyzing 10X 5′ with scCoVseq or sequencing 10X 5′ with extended R1 sequencing improved the correlation of single cell data with bulk RNA-Seq data. Combining 10X 5′ extended R1 with scCoVseq maximized quantification of sgmRNA UMIs compared to 10X 5′ or 10X 3′. Thus while analyzing 10X 5′ with scCoVseq or sequencing with extended R1 improved the confidence of sgmRNA quantification, the combination of 10X 5′ with extended R1 sequencing and scCoVseq also maximized the number of sgmRNAs detected per cell.

Our method enables the comparison of sgmRNA expression dynamics at single cell resolution. We observed that SARS-CoV-2 viral gene expression is highly correlated across cells, which suggests that across increasing levels of viral RNA, the relative proportions of viral RNA are maintained. This differs significantly from other viruses such as influenza (64, 65) and HSV (66) and CMV (67). These differences are likely due to differences in viral genome structure resulting in differences in regulation of viral gene expression. For example, the segmented nature of influenza virus genomes can result in virions missing gene segments, which partially explains increased heterogeneity in viral gene expression in influenza-infected cells (65). Future studies of coronavirus gene expression dynamics may be particularly relevant for comparing viral gene expression between different cell types, coronaviruses, or between SARS-CoV-2 variants of interest, which have been described to have different kinetics of sgmRNA expression(68). This approach could be extended to any coronavirus or nidovirus, including future emergent novel coronaviruses.

Finally, our method can be used to examine differential junction site usage within single cells (**Supplemental Figure 5**). Several groups have identified TRS-independent SARS-CoV-2 sgmRNAs(12, 13, 15), the significance of which remain unknown. It is possible that changes in junction site usage between cell types or during the course of infection may play a role in pathogenesis.

One technical advantage of our method is that it does not require modification of “out of the box” 10X library preparation steps. It therefore does not require technical optimization and is readily available to other researchers. Other methods such as virus-inclusive scRNA-Seq (69– 71), which require technical modifications or instrumentation may be less feasible for BSL-3 pathogens such as SARS-CoV-2.

It should be noted that there are some limitations to our study. With our dataset, we are unable to know the “ground truth” infection status of a cell processed for scRNA-Seq, and therefore we cannot assess the true accuracy of our method to classify infected cells. An additional limitation of our method is that quantification of viral genes with scCoVseq is dependent on accurate annotation of viral RNAs. We derived our annotation based on published empirically-defined TRS-dependent RNAs(12), but this does not preclude the existence of other viral RNAs at time points or in cell types not studied. Importantly, we explicitly exclude TRS-independent RNAs from our analyses. Methods such as STARsolo(61) or sequencing 10X libraries with long-read sequencing(65) may allow for detection and quantification of viral RNAs without reference annotation and irrespective of TRSs.

## Acknowledgments

This work was supported in part by NIH grants R21 AI149180, R01 AI151029, and U01 AI150748. P.C. was supported by the Mount Sinai Medical Scientist Training Program T32 GM007280. P.C, R.S.P., and E.J.D. were supported by the Viral Host Pathogenesis Training Grant T32 AI07647. This work was partially supported by: CRIPT (Center for Research on Influenza Pathogenesis and Transmission), a NIAID funded Center of Excellence for Influenza Research and Response (CEIRR, contract # 75N93021C00014), NIAID grants U19 AI142733 and U19 AI135972 to A.G-S, NCI Seronet grant U54 CA260560 to A.G-S., and the JPB and OPP foundations and an anonymous philanthropic donor to A.G-S.

We thank Randy Albrecht for BSL3 facility management and support. We also thank Thomas Moran, Center for Therapeutic Antibody Discovery at the Icahn School of Medicine at Mount Sinai, for kindly providing anti-SARS-CoV Nucleocapsid antibody. We thank Michael A Schotsaert for flow cytometry support. This work was supported in part through the computational resources and staff expertise provided by Scientific Computing at the Icahn School of Medicine at Mount Sinai. Research reported in this paper was supported by the Office of Research Infrastructure of the National Institutes of Health under award number S10OD026880. The content is solely the responsibility of the authors and does not necessarily represent the official views of the National Institutes of Health. Microscopy was performed at the Microscopy CoRE at the Icahn School of Medicine at Mount Sinai. NovaSeq sequencing, bulk RNA isolation, and SMARTseq library preparation was performed at the Mount Sinai Genomics Core Facility.

## Author Contributions

Conceptualization: P.C., B.R.R.

Data Curation: P.C., B.R.R.

Formal Analysis: P.C., B.R.R.

Funding Acquisition: A.G.S., B.R.R.

Investigation: P.C., E.J.D., E.B., T.K., A.C, O.D., L.M.

Methodology: P.C., B.R.R.

Project Administration: B.R.R.

Resources: A.G.S, B.R.R.

Software: P.C., R.S.P., T.B., B.R.R.,

Supervision: A.G.S., B.R.R.

Validation: P.C., R.S.P., E.J.D., O.D., T.B., E.B.,

Visualization: P.C., B.R.R.

Writing – original draft: P.C., B.R.R.

Writing – review & editing: P.C., E.J.D., O.D., R.S.P., E.B., T.B., T.K., A.C., L.M., A.G.S, B.R.R.

## References

1. Dong E, Du H, Gardner L. 2020. An interactive web-based dashboard to track COVID-19 in real time. Lancet Infect Dis 20:533–534.

2. Zhu N, Zhang D, Wang W, Li X, Yang B, Song J, Zhao X, Huang B, Shi W, Lu R, Niu P, Zhan F, Ma X, Wang D, Xu W, Wu G, Gao GF, Tan W, Investigating CNC, Team R. 2020. A novel coronavirus from patients with pneumonia in china, 2019. N Engl J Med 382:727–733.

3. Carvalho T, Krammer F, Iwasaki A. 2021. The first 12 months of COVID-19: a timeline of immunological insights. 4. Nat Rev Immunol 21:245–256.

4. Gordon DE, Jang GM, Bouhaddou M, Xu J, Obernier K, White KM, O’Meara MJ, Rezelj VV, Guo JZ, Swaney DL, Tummino TA, Huettenhain R, Kaake RM, Richards AL, Tutuncuoglu B, Foussard H, Batra J, Haas K, Modak M, Kim M, Haas P, Polacco BJ, Braberg H, Fabius JM, Eckhardt M, Soucheray M, Bennett MJ, Cakir M, McGregor MJ, Li Q, Meyer B, Roesch F, Vallet T, Mac Kain A, Miorin L, Moreno E, Naing ZZC, Zhou Y, Peng S, Shi Y, Zhang Z, Shen W, Kirby IT, Melnyk JE, Chorba JS, Lou K, Dai SA, Barrio-Hernandez I, Memon D, Hernandez-Armenta C, Lyu J, Mathy CJP, Perica T, Pilla KB, Ganesan SJ, Saltzberg DJ, Rakesh R, Liu X, Rosenthal SB, Calviello L, Venkataramanan S, Liboy-Lugo J, Lin Y, Huang X-P, Liu Y, Wankowicz SA, Bohn M, Safari M, Ugur FS, Koh C, Savar NS, Tran QD, Shengjuler D, Fletcher SJ, O’Neal MC, Cai Y, Chang JCJ, Broadhurst DJ, Klippsten S, Sharp PP, Wenzell NA, Kuzuoglu D, Wang H-Y, Trenker R, Young JM, Cavero DA, Hiatt J, Roth TL, Rathore U, Subramanian A, Noack J, Hubert M, Stroud RM, Frankel AD, Rosenberg OS, Verba KA, Agard DA, Ott M, Emerman M, Jura N, von Zastrow M, Verdin E, Ashworth A, Schwartz O, d’Enfert C, Mukherjee S, Jacobson M, Malik HS, Fujimori DG, Ideker T, Craik CS, Floor SN, Fraser JS, Gross JD, Sali A, Roth BL, Ruggero D, Taunton J, Kortemme T, Beltrao P, Vignuzzi M, García-Sastre A, Shokat KM, Shoichet BK, Krogan NJ. 2020. A SARS-CoV-2 protein interaction map reveals targets for drug repurposing. Nature https://doi.org/10.1038/s41586-020-2286-9.

5. Shen B, Yi X, Sun Y, Bi X, Du J, Zhang C, Quan S, Zhang F, Sun R, Qian L, Ge W, Liu W, Liang S, Chen H, Zhang Y, Li J, Xu J, He Z, Chen B, Wang J, Yan H, Zheng Y, Wang D, Zhu J, Kong Z, Kang Z, Liang X, Ding X, Ruan G, Xiang N, Cai X, Gao H, Li L, Li S, Xiao Q, Lu T, Zhu Y, Liu H, Chen H, Guo T. 2020. Proteomic and metabolomic characterization of COVID-19 patient sera. Cell https://doi.org/10.1016/j.cell.2020.05.032.

6. Bojkova D, Klann K, Koch B, Widera M, Krause D, Ciesek S, Cinatl J, Münch C. 2020. SARS-CoV-2 infected host cell proteomics reveal potential therapy targets https://doi.org/10.21203/rs.3.rs-17218/v1.

7. Finkel Y, Mizrahi O, Nachshon A, Weingarten-Gabbay S, Morgenstern D, Yahalom-Ronen Y, Tamir H, Achdout H, Stein D, Israeli O, Beth-Din A, Melamed S, Weiss S, Israely T, Paran N, Schwartz M, Stern-Ginossar N. 2020. The coding capacity of SARS-CoV-2. Nature 1–9.

8. Bouhaddou M, Memon D, Meyer B, White KM, Rezelj VV, Marrero MC, Polacco BJ, Melnyk JE, Ulferts S, Kaake RM, Batra J, Richards AL, Stevenson E, Gordon DE, Rojc A, Obernier K, Fabius JM, Soucheray M, Miorin L, Moreno E, Koh C, Tran QD, Hardy A, Robinot R, Vallet T, Nilsson-Payant BE, Hernandez-Armenta C, Dunham A, Weigang S, Knerr J, Modak M, Quintero D, Zhou Y, Dugourd A, Valdeolivas A, Patil T, Li Q, Hüttenhain R, Cakir M, Muralidharan M, Kim M, Jang G, Tutuncuoglu B, Hiatt J, Guo JZ, Xu J, Bouhaddou S, Mathy CJP, Gaulton A, Manners EJ, Félix E, Shi Y, Goff M, Lim JK, McBride T, O’Neal MC, Cai Y, Chang JCJ, Broadhurst DJ, Klippsten S, Wit ED, Leach AR, Kortemme T, Shoichet B, Ott M, Saez-Rodriguez J, tenOever BR, Mullins D, Fischer ER, Kochs G, Grosse R, García-Sastre A, Vignuzzi M, Johnson JR, Shokat KM, Swaney DL, Beltrao P, Krogan NJ. 2020. The Global Phosphorylation Landscape of SARS-CoV-2 Infection. Cell 0.

9. Schmidt N, Lareau CA, Keshishian H, Ganskih S, Schneider C, Hennig T, Melanson R, Werner S, Wei Y, Zimmer M, Ade J, Kirschner L, Zielinski S, Dölken L, Lander ES, Caliskan N, Fischer U, Vogel J, Carr SA, Bodem J, Munschauer M. 2021. The SARS-CoV-2 RNA– protein interactome in infected human cells. 3. Nat Microbiol 6:339–353.

10. Flynn RA, Belk JA, Qi Y, Yasumoto Y, Wei J, Alfajaro MM, Shi Q, Mumbach MR, Limaye A, DeWeirdt PC, Schmitz CO, Parker KR, Woo E, Chang HY, Horvath TL, Carette JE, Bertozzi CR, Wilen CB, Satpathy AT. 2021. Discovery and functional interrogation of SARS-CoV-2 RNA-host protein interactions. Cell 184:2394-2411.e16.

11. Blanco-Melo D, Nilsson-Payant BE, Liu W-C, Uhl S, Hoagland D, Møller R, Jordan TX, Oishi K, Panis M, Sachs D, Wang TT, Schwartz RE, Lim JK, Albrecht RA, tenOever BR. 2020. Imbalanced Host Response to SARS-CoV-2 Drives Development of COVID-19. Cell 181:1036-1045.e9.

12. Kim D, Lee J-Y, Yang J-S, Kim JW, Kim VN, Chang H. 2020. The architecture of SARS-CoV-2 transcriptome. Cell 181:914-921.e10.

13. Wang D, Jiang A, Feng J, Li G, Guo D, Sajid M, Wu K, Zhang Q, Ponty Y, Will S, Liu F, Yu X, Li S, Liu Q, Yang X-L, Guo M, Li X, Chen M, Shi Z-L, Lan K, Chen Y, Zhou Y. 2021. The SARS-CoV-2 Subgenome Landscape and its Novel Regulatory Features. Mol Cell 0.

14. Ziv O, Price J, Shalamova L, Kamenova T, Goodfellow I, Weber F, Miska EA. 2020. The short- and long-range RNA-RNA Interactome of SARS-CoV-2. Mol Cell https://doi.org/10.1016/j.molcel.2020.11.004.

15. Chang JJ-Y, Rawlinson D, Pitt ME, Taiaroa G, Gleeson J, Zhou C, Mordant FL, Paoli-Iseppi RD, Caly L, Purcell DFJ, Stinear TP, Londrigan SL, Clark MB, Williamson DA, Subbarao K, Coin LJM. 2021. Transcriptional and epi-transcriptional dynamics of SARS-CoV-2 during cellular infection. Cell Rep 0.

16. Bost P, Giladi A, Liu Y, Bendjelal Y, Xu G, David E, Blecher-Gonen R, Cohen M, Medaglia C, Li H, Deczkowska A, Zhang S, Schwikowski B, Zhang Z, Amit I. 2020. Host-viral infection maps reveal signatures of severe COVID-19 patients. Cell https://doi.org/10.1016/j.cell.2020.05.006.

17. Sungnak W, Huang N, Bécavin C, Berg M, Queen R, Litvinukova M, Talavera-López C, Maatz H, Reichart D, Sampaziotis F, Worlock KB, Yoshida M, Barnes JL, Network HLB. 2020. SARS-CoV-2 entry factors are highly expressed in nasal epithelial cells together with innate immune genes. Nat Med 26:681–687.

18. Lukassen S, Chua RL, Trefzer T, Kahn NC, Schneider MA, Muley T, Winter H, Meister M, Veith C, Boots AW, Hennig BP, Kreuter M, Conrad C, Eils R. 2020. SARS-CoV-2 receptor ACE2 and TMPRSS2 are primarily expressed in bronchial transient secretory cells. EMBO J 39:e105114.

19. Fodoulian L, Tuberosa J, Rossier D, Landis B, Carleton A, Rodriguez I. 2020. SARS-CoV-2 receptor and entry genes are expressed by sustentacular cells in the human olfactory neuroepithelium. BioRxiv https://doi.org/10.1101/2020.03.31.013268.

20. Qi F, Qian S, Zhang S, Zhang Z. 2020. Single cell RNA sequencing of 13 human tissues identify cell types and receptors of human coronaviruses. Biochem Biophys Res Commun 526:135–140.

21. Ravindra NG, Alfajaro MM, Gasque V, Huston NC, Wan H, Szigeti-Buck K, Yasumoto Y, Greaney AM, Habet V, Chow RD, Chen JS, Wei J, Filler RB, Wang B, Wang G, Niklason LE, Montgomery RR, Eisenbarth SC, Chen S, Williams A, Iwasaki A, Horvath TL, Foxman EF, Pierce RW, Pyle AM, Dijk D van, Wilen CB. 2021. Single-cell longitudinal analysis of SARS-CoV-2 infection in human airway epithelium identifies target cells, alterations in gene expression, and cell state changes. PLOS Biol 19:e3001143.

22. Muus C, Luecken MD, Eraslan G, Sikkema L, Waghray A, Heimberg G, Kobayashi Y, Vaishnav ED, Subramanian A, Smillie C, Jagadeesh KA, Duong ET, Fiskin E, Torlai Triglia E, Ansari M, Cai P, Lin B, Buchanan J, Chen S, Shu J, Haber AL, Chung H, Montoro DT, Adams T, Aliee H, Allon SJ, Andrusivova Z, Angelidis I, Ashenberg O, Bassler K, Bécavin C, Benhar I, Bergenstråhle J, Bergenstråhle L, Bolt L, Braun E, Bui LT, Callori S, Chaffin M, Chichelnitskiy E, Chiou J, Conlon TM, Cuoco MS, Cuomo ASE, Deprez M, Duclos G, Fine D, Fischer DS, Ghazanfar S, Gillich A, Giotti B, Gould J, Guo M, Gutierrez AJ, Habermann AC, Harvey T, He P, Hou X, Hu L, Hu Y, Jaiswal A, Ji L, Jiang P, Kapellos TS, Kuo CS, Larsson L, Leney-Greene MA, Lim K, Litviňuková M, Ludwig LS, Lukassen S, Luo W, Maatz H, Madissoon E, Mamanova L, Manakongtreecheep K, Leroy S, Mayr CH, Mbano IM, McAdams AM, Nabhan AN, Nyquist SK, Penland L, Poirion OB, Poli S, Qi C, Queen R, Reichart D, Rosas I, Schupp JC, Shea CV, Shi X, Sinha R, Sit RV, Slowikowski K, Slyper M, Smith NP, Sountoulidis A, Strunz M, Sullivan TB, Sun D, Talavera-López C, Tan P, Tantivit J, Travaglini KJ, Tucker NR, Vernon KA, Wadsworth MH, Waldman J, Wang X, Xu K, Yan W, Zhao W, Ziegler CGK. 2021. Single-cell meta-analysis of SARS-CoV-2 entry genes across tissues and demographics. 3. Nat Med 27:546–559.

23. Bieberich F, Vazquez-Lombardi R, Yermanos A, Ehling RA, Mason DM, Wagner B, Kapetanovic E, Roberto RBD, Weber CR, Savic M, Rudolf F, Reddy ST. 2021. A single-cell atlas of lymphocyte adaptive immune repertoires and transcriptomes reveals age-related differences in convalescent COVID-19 patients. bioRxiv 2021.02.12.430907.

24. Wilk AJ, Rustagi A, Zhao NQ, Roque J, Martínez-Colón GJ, McKechnie JL, Ivison GT, Ranganath T, Vergara R, Hollis T, Simpson LJ, Grant P, Subramanian A, Rogers AJ, Blish CA. 2020. A single-cell atlas of the peripheral immune response in patients with severe COVID-19. 7. Nat Med 26:1070–1076.

25. Yao C, Bora SA, Parimon T, Zaman T, Friedman OA, Palatinus JA, Surapaneni NS, Matusov YP, Chiang GC, Kassar AG, Patel N, Green CER, Aziz AW, Suri H, Suda J, Lopez AA, Martins GA, Stripp BR, Gharib SA, Goodridge HS, Chen P. 2021. Cell-Type-Specific Immune Dysregulation in Severely Ill COVID-19 Patients. Cell Rep 34.

26. MacDonald L, Otto TD, Elmesmari A, Tolusso B, Somma D, McSharry C, Gremese E, McInnes IB, Alivernini S, Kurowska-Stolarska M. 2020. COVID-19 and Rheumatoid Arthritis share myeloid pathogenic and resolving pathways. bioRxiv 2020.07.26.221572.

27. Schreibing F, Hannani M, Ticconi F, Fewings E, Nagai JS, Begemann M, Kuppe C, Kurth I, Kranz J, Frank D, Anslinger TM, Ziegler P, Kraus T, Enczmann J, Balz V, Windhofer F, Balfanz P, Kurts C, Marx G, Marx N, Dreher M, Schneider RK, Saez-Rodriguez J, Filho IGC, Kramann R. 2021. Dissecting CD8+ T cell pathology of severe SARS-CoV-2 infection by single-cell epitope mapping. bioRxiv 2021.03.03.432690.

28. Wen W, Su W, Tang H, Le W, Zhang X, Zheng Y, Liu X, Xie L, Li J, Ye J, Dong L, Cui X, Miao Y, Wang D, Dong J, Xiao C, Chen W, Wang H. 2020. Immune cell profiling of COVID-19 patients in the recovery stage by single-cell sequencing. Cell Discov 6:31.

29. Lee JS, Park S, Jeong HW, Ahn JY, Choi SJ, Lee H, Choi B, Nam SK, Sa M, Kwon J-S, Jeong SJ, Lee HK, Park SH, Park S-H, Choi JY, Kim S-H, Jung I, Shin E-C. 2020. Immunophenotyping of COVID-19 and influenza highlights the role of type I interferons in development of severe COVID-19. Sci Immunol 5.

30. Wang F-S, Zhang J-Y, Wang X, Xing X, Xu Z, Zhang C, Song J-W, Fan X, Xia P, Fu J-L, Wang S-Y, Xu R-N, Dai X-P, Shi L, Huang L, Jiang T-J, Shi M, Zhang Y, Zumla A, Maeurer M, Bai F. 2020. Single-cell landscape of immunological responses in COVID-19 patients. bioRxiv 2020.07.23.217703.

31. Zhang J-Y, Wang X-M, Xing X, Xu Z, Zhang C, Song J-W, Fan X, Xia P, Fu J-L, Wang S-Y, Xu R-N, Dai X-P, Shi L, Huang L, Jiang T-J, Shi M, Zhang Y, Zumla A, Maeurer M, Bai F, Wang F-S. 2020. Single-cell landscape of immunological responses in patients with COVID-19. 9. Nat Immunol 21:1107–1118.

32. Kalfaoglu B, Almeida-Santos J, Tye CA, Satou Y, Ono M. 2020. T-cell hyperactivation and paralysis in severe COVID-19 infection revealed by single-cell analysis. BioRxiv https://doi.org/10.1101/2020.05.26.115923.

33. Wei L, Ming S, Zou B, Wu Y, Hong Z, Li Z, Zheng X, Huang M, Luo L, Liang J, Wen X, Chen T, Liang Q, Kuang L, Shan H, Huang X. 2020. Viral invasion and type I interferon response characterize the immunophenotypes during COVID-19 infection. Electron J https://doi.org/10.2139/ssrn.3555695.

34. Wyler E, Mösbauer K, Franke V, Diag A, Gottula LT, Arsie R, Klironomos F, Koppstein D, Ayoub S, Buccitelli C, Richter A, Legnini I, Ivanov A, Mari T, Del Giudice S, Papies JP, Müller MA, Niemeyer D, Selbach M, Akalin A, Rajewsky N, Drosten C, Landthaler M. 2020. Bulk and single-cell gene expression profiling of SARS-CoV-2 infected human cell lines identifies molecular targets for therapeutic intervention. BioRxiv https://doi.org/10.1101/2020.05.05.079194.

35. Miorin L, Kehrer T, Sanchez-Aparicio MT, Zhang K, Cohen P, Patel RS, Cupic A, Makio T, Mei M, Moreno E, Danziger O, White KM, Rathnasinghe R, Uccellini M, Gao S, Aydillo T, Mena I, Yin X, Martin-Sancho L, Krogan NJ, Chanda SK, Schotsaert M, Wozniak RW, Ren Y, Rosenberg BR, Fontoura BMA, García-Sastre A. 2020. SARS-CoV-2 Orf6 hijacks Nup98 to block STAT nuclear import and antagonize interferon signaling. Proc Natl Acad Sci 117:28344–28354.

36. Melms JC, Biermann J, Huang H, Wang Y, Nair A, Tagore S, Katsyv I, Rendeiro AF, Amin AD, Schapiro D, Frangieh CJ, Luoma AM, Filliol A, Fang Y, Ravichandran H, Clausi MG, Alba GA, Rogava M, Chen SW, Ho P, Montoro DT, Kornberg AE, Han AS, Bakhoum MF, Anandasabapathy N, Suárez-Fariñas M, Bakhoum SF, Bram Y, Borczuk A, Guo XV, Lefkowitch JH, Marboe C, Lagana SM, Del Portillo A, Zorn E, Markowitz GS, Schwabe RF, Schwartz RE, Elemento O, Saqi A, Hibshoosh H, Que J, Izar B. 2021. A molecular single-cell lung atlas of lethal COVID-19. Nature 1–6.

37. Delorey TM, Ziegler CGK, Heimberg G, Normand R, Yang Y, Segerstolpe Å, Abbondanza D, Fleming SJ, Subramanian A, Montoro DT, Jagadeesh KA, Dey KK, Sen P, Slyper M, Pita-Juárez YH, Phillips D, Biermann J, Bloom-Ackermann Z, Barkas N, Ganna A, Gomez J, Melms JC, Katsyv I, Normandin E, Naderi P, Popov YV, Raju SS, Niezen S, Tsai LT-Y, Siddle KJ, Sud M, Tran VM, Vellarikkal SK, Wang Y, Amir-Zilberstein L, Atri DS, Beechem J, Brook OR, Chen J, Divakar P, Dorceus P, Engreitz JM, Essene A, Fitzgerald DM, Fropf R, Gazal S, Gould J, Grzyb J, Harvey T, Hecht J, Hether T, Jané-Valbuena J, Leney-Greene M, Ma H, McCabe C, McLoughlin DE, Miller EM, Muus C, Niemi M, Padera R, Pan L, Pant D, Pe’er C, Pfiffner-Borges J, Pinto CJ, Plaisted J, Reeves J, Ross M, Rudy M, Rueckert EH, Siciliano M, Sturm A, Todres E, Waghray A, Warren S, Zhang S, Zollinger DR, Cosimi L, Gupta RM, Hacohen N, Hibshoosh H, Hide W, Price AL, Rajagopal J, Tata PR, Riedel S, Szabo G, Tickle TL, Ellinor PT, Hung D, Sabeti PC, Novak R, Rogers R, Ingber DE, Jiang ZG, Juric D, Babadi M, Farhi SL, Izar B, Stone JR, Vlachos IS, Solomon IH, Ashenberg O, Porter CBM, Li B, Shalek AK, Villani A-C, Rozenblatt-Rosen O, Regev A. 2021. COVID-19 tissue atlases reveal SARS-CoV-2 pathology and cellular targets. Nature 1–8.

38. Harrison AG, Lin T, Wang P. 2020. Mechanisms of SARS-CoV-2 Transmission and Pathogenesis. Trends Immunol 41:1100–1115.

39. V’kovski P, Kratzel A, Steiner S, Stalder H, Thiel V. 2020. Coronavirus biology and replication: implications for SARS-CoV-2. 3. Nat Rev Microbiol 1–16.

40. Sola I, Almazán F, Zúñiga S, Enjuanes L. 2015. Continuous and discontinuous RNA synthesis in coronaviruses. Annu Rev Virol 2:265–288.

41. Perlman S, Masters PS. 2020. Coronaviridae: The Viruses and Their Replication, p. 411–442. In Howley, PM, Knipe, DM, Whelan, S (eds.), Fields Virology: Emerging Viruses, 7th ed. Wolters Kluwer Health/lippincott Williams & Wilkins, Philadelphia, PA.

42. Parker MD, Lindsey BB, Leary S, Gaudieri S, Chopra A, Wyles M, Angyal A, Green LR, Parsons P, Tucker RM, Brown R, Groves D, Johnson K, Carrilero L, Heffer J, Partridge DG, Evans C, Raza M, Keeley AJ, Smith N, Filipe ADS, Shepherd JG, Davis C, Bennett S, Sreenu VB, Kohl A, Aranday-Cortes E, Tong L, Nichols J, Thomson EC, Consortium TC-19 GU (COG-U, Wang D, Mallal S, Silva TI de. 2021. Subgenomic RNA identification in SARS-CoV-2 genomic sequencing data. Genome Res 31:645–658.

43. Daniloski Z, Jordan TX, Wessels H-H, Hoagland DA, Kasela S, Legut M, Maniatis S, Mimitou EP, Lu L, Geller E, Danziger O, Rosenberg BR, Phatnani H, Smibert P, Lappalainen T, tenOever BR, Sanjana NE. 2021. Identification of Required Host Factors for SARS-CoV-2 Infection in Human Cells. Cell 184:92-105.e16.

44. Miorin L, Mire CE, Ranjbar S, Hume AJ, Huang J, Crossland NA, White KM, Laporte M, Kehrer T, Haridas V, Moreno E, Nambu A, Jangra S, Cupic A, Dejosez M, Abo KA, Tseng AE, Werder RB, Rathnasinghe R, Mutetwa T, Ramos I, Aja JS de, Rivas CG de A, Schotsaert M, Corley RB, Falvo JV, Fernandez-Sesma A, Kim C, Rossignol J-F, Wilson AA, Zwaka T, Kotton DN, Mühlberger E, García-Sastre A, Goldfeld AE. 2022. The oral drug nitazoxanide restricts SARS-CoV-2 infection and attenuates disease pathogenesis in Syrian hamsters. bioRxiv https://doi.org/10.1101/2022.02.08.479634.

45. Kehrer T, Cupic A, Ye C, Yildiz S, Bouhhadou M, Crossland NA, Barrall E, Cohen P, Tseng A, Çağatay T, Rathnasinghe R, Flores D, Jangra S, Alam F, Mena N, Aslam S, Saqi A, Marin A, Rutkowska M, Ummadi MR, Pisanelli G, Richardson RB, Veit EC, Fabius JM, Soucheray M, Polacco BJ, Evans MJ, Swaney DL, Gonzalez-Reiche AS, Sordillo EM, Bakel H van, Simon V, Zuliani-Alvarez L, Fontoura BMA, Rosenberg BR, Krogan NJ, Martinez-Sobrido L, García-Sastre A, Miorin L. 2022. Impact of SARS-CoV-2 ORF6 and its variant polymorphisms on host responses and viral pathogenesis. bioRxiv https://doi.org/10.1101/2022.10.18.512708.

46. Gonzalez-Reiche AS, Hernandez MM, Sullivan MJ, Ciferri B, Alshammary H, Obla A, Fabre S, Kleiner G, Polanco J, Khan Z, Alburquerque B, van de Guchte A, Dutta J, Francoeur N, Melo BS, Oussenko I, Deikus G, Soto J, Sridhar SH, Wang Y-C, Twyman K, Kasarskis A, Altman DR, Smith M, Sebra R, Aberg J, Krammer F, García-Sastre A, Luksza M, Patel G, Paniz-Mondolfi A, Gitman M, Sordillo EM, Simon V, van Bakel H. 2020. Introductions and early spread of SARS-CoV-2 in the New York City area. Science https://doi.org/10.1126/science.abc1917.

47. Escalera A, Gonzalez-Reiche AS, Aslam S, Mena I, Laporte M, Pearl RL, Fossati A, Rathnasinghe R, Alshammary H, van de Guchte A, Farrugia K, Qin Y, Bouhaddou M, Kehrer T, Zuliani-Alvarez L, Meekins DA, Balaraman V, McDowell C, Richt JA, Bajic G, Sordillo EM, Dejosez M, Zwaka TP, Krogan NJ, Simon V, Albrecht RA, van Bakel H, García-Sastre A, Aydillo T. 2022. Mutations in SARS-CoV-2 variants of concern link to increased spike cleavage and virus transmission. Cell Host Microbe 30:373-387.e7.

48. Macosko EZ, Basu A, Satija R, Nemesh J, Shekhar K, Goldman M, Tirosh I, Bialas AR, Kamitaki N, Martersteck EM, Trombetta JJ, Weitz DA, Sanes JR, Shalek AK, Regev A, McCarroll SA. 2015. Highly parallel genome-wide expression profiling of individual cells using nanoliter droplets. Cell 161:1202–1214.

49. Butler A, Hoffman P, Smibert P, Papalexi E, Satija R. 2018. Integrating single-cell transcriptomic data across different conditions, technologies, and species. Nat Biotechnol 36:411–420.

50. Hafemeister C, Satija R. 2019. Normalization and variance stabilization of single-cell RNA-seq data using regularized negative binomial regression. Genome Biol 20:296.

51. Li H. 2021. lh3/seqtk. C.

52. Fernandes JD, Hinrichs AS, Clawson H, Navarro Gonzalez J, Lee BT, Nassar LR, Raney BJ, Rosenbloom KR, Nerli S, Rao A, Schmelter D, Zweig AS, Lowe TM, Ares M, Corbet-Detig R, Kent WJ, Haussler D, Haeussler M. 2020. The UCSC SARS-CoV-2 genome browser. BioRxiv https://doi.org/10.1101/2020.05.04.075945.

53. Danecek P, Bonfield JK, Liddle J, Marshall J, Ohan V, Pollard MO, Whitwham A, Keane T, McCarthy SA, Davies RM, Li H. 2021. Twelve years of SAMtools and BCFtools. GigaScience 10:giab008.

54. Smith T, Heger A, Sudbery I. 2017. UMI-tools: modeling sequencing errors in Unique Molecular Identifiers to improve quantification accuracy. Genome Res 27:491–499.

55. 2021. Sparse and Dense Matrix Classes and Methods [R package Matrix version 1.3-4]. Comprehensive R Archive Network (CRAN). https://CRAN.R-project.org/package=Matrix. Retrieved 3 June 2021.

56. 2020. Read Rectangular Text Data [R package readr version 1.4.0]. Comprehensive R Archive Network (CRAN). https://CRAN.R-project.org/package=readr. Retrieved 3 June 2021.

57. Garrido-Martín D, Palumbo E, Guigó R, Breschi A. 2018. ggsashimi: Sashimi plot revised for browser- and annotation-independent splicing visualization. PLOS Comput Biol 14:e1006360.

58. Maechler M, original) PR (Fortran, original) AS (S, original) MH (S, Hornik [trl K, maintenance(1999-2000)) ctb] (port to R, Studer M, Roudier P, Gonzalez J, Kozlowski K, pam()) ES (fastpam options for, Murphy (volume.ellipsoid({d >= 3})) K. 2021. cluster: “Finding Groups in Data”: Cluster Analysis Extended Rousseeuw et al. (2.1.2).

59. Robinson MD, McCarthy DJ, Smyth GK. 2010. edgeR: a Bioconductor package for differential expression analysis of digital gene expression data. Bioinformatics 26:139–140.

60. Soneson C, Robinson MD. 2018. Bias, robustness and scalability in single-cell differential expression analysis. 4. Nat Methods 15:255–261.

61. Kaminow B, Yunusov D, Dobin A. 2021. STARsolo: accurate, fast and versatile mapping/quantification of single-cell and single-nucleus RNA-seq data. bioRxiv 2021.05.05.442755.

62. McQuin C, Goodman A, Chernyshev V, Kamentsky L, Cimini BA, Karhohs KW, Doan M, Ding L, Rafelski SM, Thirstrup D, Wiegraebe W, Singh S, Becker T, Caicedo JC, Carpenter AE. 2018. CellProfiler 3.0: Next-generation image processing for biology. PLOS Biol 16:e2005970.

63. Emeny JM, Morgan MJY 1979. 1979. Regulation of the Interferon System: Evidence that Vero Cells have a Genetic Defect in Interferon Production. J Gen Virol 43:247–252.

64. Russell AB, Trapnell C, Bloom JD. 2018. Extreme heterogeneity of influenza virus infection in single cells. eLife 7:e32303.

65. Russell AB, Elshina E, Kowalsky JR, Te Velthuis Ajw, Bloom JD. 2019. Single-cell virus sequencing of influenza infections that trigger innate immunity. J Virol 93.

66. Drayman N, Patel P, Vistain L, Tay S. 2019. HSV-1 single-cell analysis reveals the activation of anti-viral and developmental programs in distinct sub-populations. eLife 8:e46339.

67. Hein MY, Weissman JS. 2022. Functional single-cell genomics of human cytomegalovirus infection. 3. Nat Biotechnol 40:391–401.

68. Parker MD, Lindsey BB, Shah DR, Hsu S, Keeley AJ, Partridge DG, Leary S, Cope A, State A, Johnson K, Ali N, Raghei R, Heffer J, Smith N, Zhang P, Gallis M, Louka SF, Whiteley M, Foulkes BH, Christou S, Wolverson P, Pohare M, Hansford SE, Green LR, Evans C, Raza M, Wang D, Gaudieri S, Mallal S, Consortium TC-19 GU (COG-U, Silva TI de. 2021. Altered Sub-Genomic RNA Expression in SARS-CoV-2 B.1.1.7 Infections. bioRxiv 2021.03.02.433156.

69. Zanini F, Pu S-Y, Bekerman E, Einav S, Quake SR. 2018. Single-cell transcriptional dynamics of flavivirus infection. eLife 7.

70. Zanini F, Robinson ML, Croote D, Sahoo MK, Sanz AM, Ortiz-Lasso E, Albornoz LL, Rosso F, Montoya JG, Goo L, Pinsky BA, Quake SR, Einav S. 2018. Virus-inclusive single-cell RNA sequencing reveals the molecular signature of progression to severe dengue. Proc Natl Acad Sci U S A 115:E12363–E12369.

71. O’Neal JT, Upadhyay AA, Wolabaugh A, Patel NB, Bosinger SE, Suthar MS. 2019. West nile virus-inclusive single-cell RNA sequencing reveals heterogeneity in the type I interferon response within single cells. J Virol 93.

