## Supplemental Figures for "Unambiguous detection of SARS-CoV-2 subgenomic mRNAs with single cell RNA sequencing"

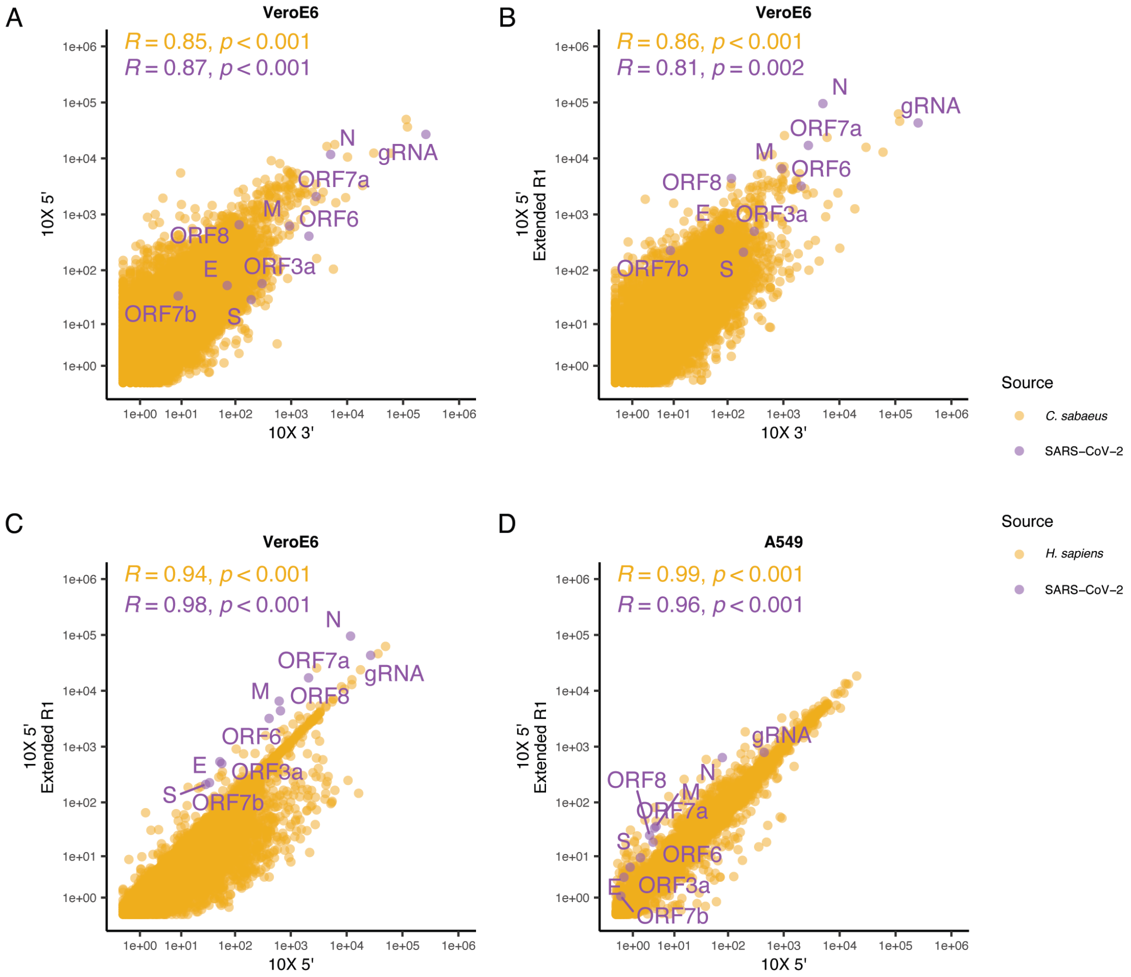


**Supplemental Figure 1:** Host (orange) and virus (purple) gene expression in Vero E6 cells (**A-C**) and ACE2-A549 cells (**D**) measured by 10X 3′, 10X 5′, and 10X 5′ with extended R1 sequencing. Each point represents the aggregate pseudobulk UMI count for a single gene assayed in the indicated assays per $10^6$ total pseudobulk UMIs. Pearson correlation coefficients and resulting p values are indicated in each plot.

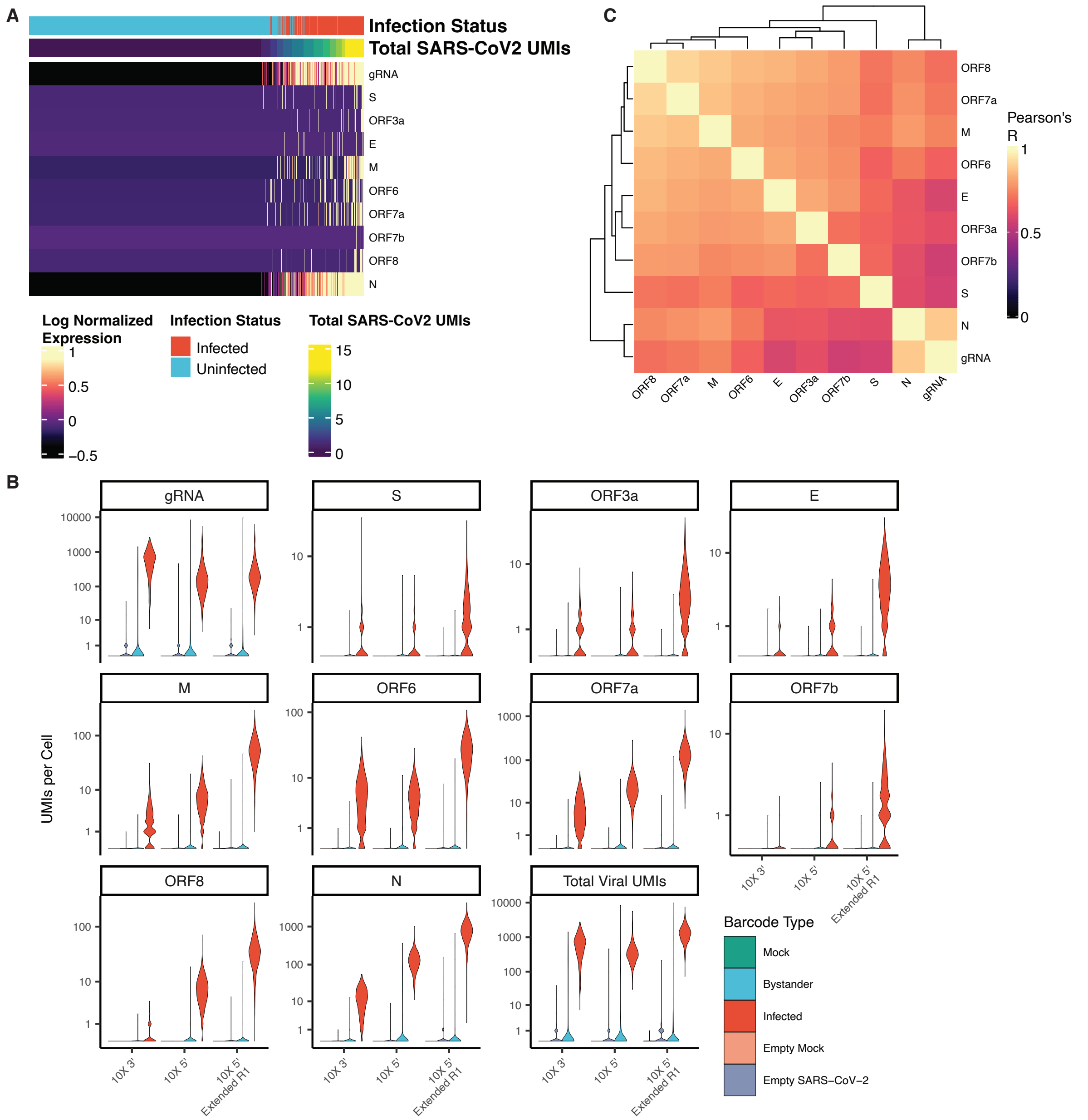


**Supplemental Figure 2: A.** Expression of viral genes in rSARS-CoV-2 ORF6 M58R infected and uninfected ACE2-A549 cells. Cells are represented as columns with the expression of each viral gene indicated by color. Expression values for each gene are “clipped” to the 95$^{th}$ percentile of log normalized expression of all genes across all plotted cells to improve visibility. Cells are ordered left to right by increasing total SARS-CoV-2 viral UMIs. Total SARS-CoV-2 viral UMI counts are “clipped” to the value of the 95$^{th}$ percentile of total viral UMIs to improve visibility. Cells are labelled above with their assigned infection status. **B.** Viral gene expression of infected and bystander Vero E6 cells as well as showing sampled empty droplets from the mock and SARS-CoV-2 treated sample. **C.** Correlation of rSARS-CoV-2 ORF6 M58R viral gene expression across ACE2-A549 cells as measured by 10X 5′ extended R1.

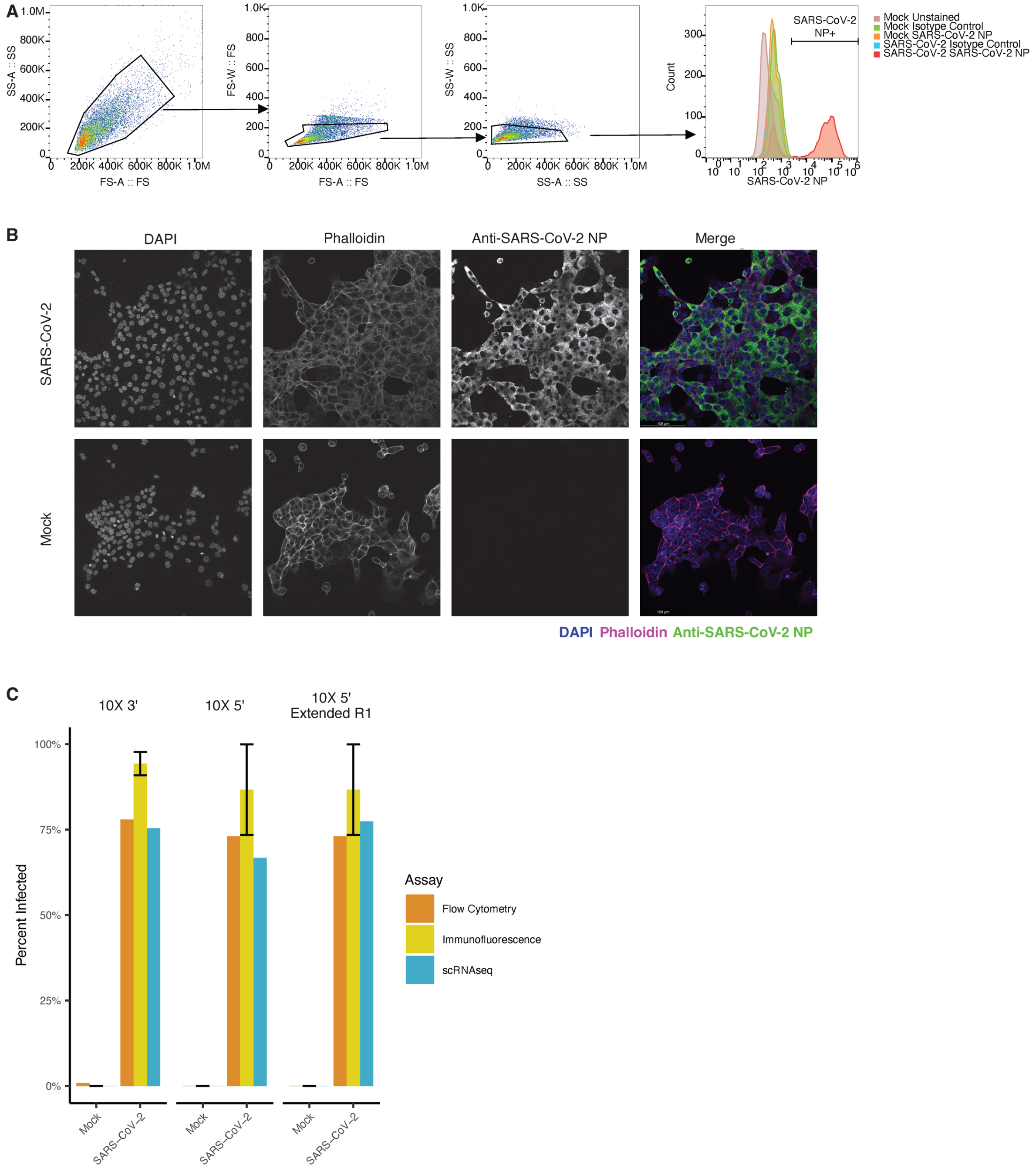


**Supplemental Figure 3**: **A.** Example flow cytometry plots illustrating gating and SARS-CoV-2 NP labeling intensities in Vero E6 infected and control samples. **B.** Representative microscopy images of Vero E6 SARS-CoV-2 NP labeling for infected and mock cells. **C.** Percent of infected cells per sample (Vero E6 cells) as measured by flow cytometry, immunofluorescence, and scRNA-Seq. Because the same sample was sequenced with 10X 5′ and 10X 5′ extended R1, flow cytometry and immunofluorescence results are duplicated for ease of visualization. Error bars for immunofluorescence indicate mean ± one standard deviation of percent infected cells based on three fields per sample.

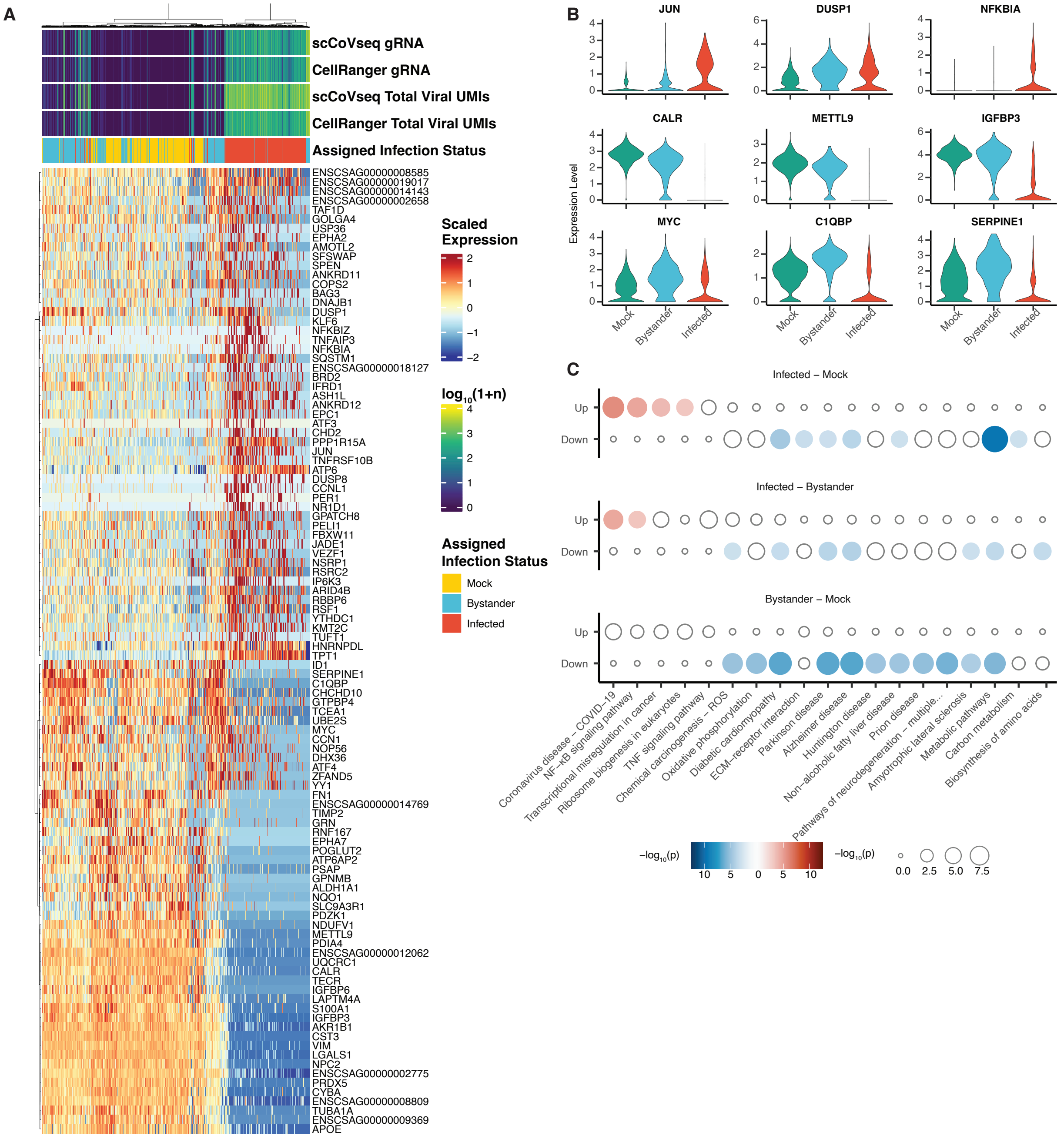


**Supplemental Figure 4: A.** Heatmap of genes differentially expressed in infected, bystander, or mock Vero E6 cells. Differential expression testing was performed on host gene expression downsampled to an equal number of UMIs/cell across cells to account for infection-induced transcriptional shutdown. Genes were selected for visualization based on false discovery rate of less than 0.05 and absolute $\log_2$ fold change of at least 1. Non-downsampled gene expression data is shown. Along the top, infection status, total viral UMIs and genomic RNA as quantified by CellRanger and scCoVseq are indicated. Cells and genes are clustered with Ward d2 clustering on Euclidean distance. **B.** Expression of select host genes per cell by infection status. Data shown are not downsampled. *Top:* genes induced in infected cells. *Middle:* genes repressed in infected cells. *Bottom:* genes upregulated in bystander cells compared to mock. **C.** KEGG pathway enrichment of genes differentially expressed in pairwise comparisons of downsampled infected, bystander, and mock Vero E6 cells. Dot size and fill indicates the $-\log_{10}$ p value of enrichment with red dots indicating enrichment in the first infection state and blue in the second infection state noted above each panel.

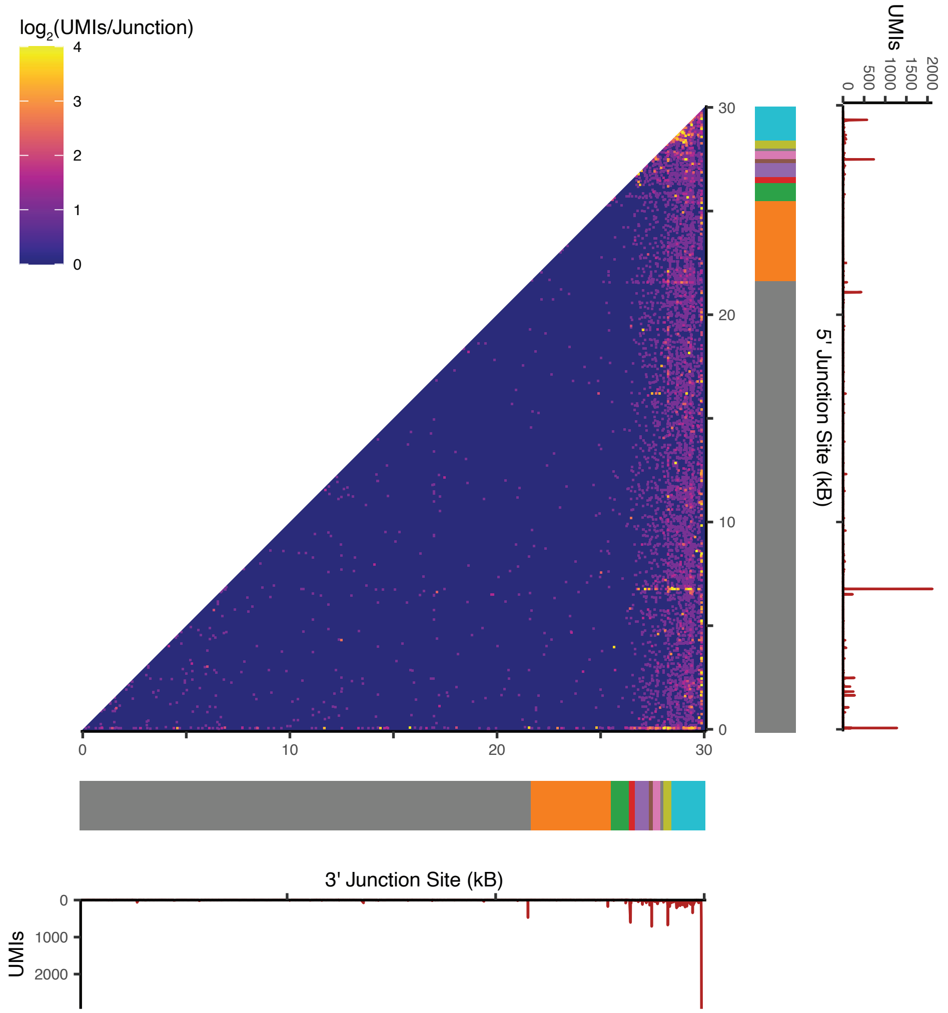


**Supplemental Figure 5:** Detection of junction sites in SARS-CoV-2 reads with 10X 5′ extended R1 (Vero E6 cells). Junction sites are represented by the 5′ start site and 3′ end site on the y and x-axis, respectively. The color indicates the $\log_2$ total UMIs/junction across all cells in the SARS-CoV-2 infected sample. Below each axis, the number of UMIs supporting a position as a junction start or end site is indicated with a density plot.
